# Nutrient excess remodels islet autonomic innervation via pancreatic schwann cells

**DOI:** 10.1101/2025.07.01.662442

**Authors:** R.F. Hampton, M. Jimenez-Gonzalez, R. Kang, A. Alvarsson, D. Espinoza, J.R.E. Carty, F. Seibold, L. Lambertini, L. Alexeyev, R. Li, K. Devarakonda, G. Lu, B. Agyapong, J. Choudhury, M. Alon, D. Scott, A. Stewart, A. Garcia-Ocana, S.A. Stanley

**Affiliations:** Diabetes, Obesity and Metabolism Institute, Icahn School of Medicine at Mount Sinai, New York, NY 10029, USA; Nash Family Department of Neuroscience and Friedman Brain Institute, Icahn School of Medicine at Mount Sinai, New York, NY 10029, USA; Department of Molecular and Cellular Endocrinology, Arthur Riggs Diabetes and Metabolism Research Institute, City of Hope Beckman Research Institute, Duarte, CA 91010, USA

## Abstract

Nutrient excess results in short- and long-term adaptations in the structure and function of metabolic organs, such as pancreatic islets. Pancreatic innervation contributes to islet architecture and function in many species, including humans, but little is known about its adaptation to over-nutrition. Here, we use 3D imaging to show that short-term high fat diet rapidly remodels islet sympathetic innervation while long-term high fat diet results in islet-specific cholinergic neuropathy. Using highly targeted, organ- and circuit-specific approaches, we demonstrate that remodeling of pancreatic innervation contributes to impaired glycemic control with short- and long-term nutrient excess. Counteracting the nutrient-induced changes in sympathetic and parasympathetic inputs to the pancreas using neuromodulation and repurposed FDA-approved approaches, we can significantly improve glycemic control even with long-term high fat diet. Finally, we identify islet Schwann cells as a source of the neurotrophic factor, S100b, contributing to rapid sympathetic hyperinnervation with short-term overnutrition. Our findings reveal novel adaptations in islet innervation contributing to nutrition-mediated metabolic dysfunction that can be ameliorated by targeting pancreatic neural circuits.

---

Food and nutrient availability can be unpredictable in the natural environment with some periods of abundance and others of limited food availability. When food is readily available, metabolic organs, such as the endocrine pancreas, show rapid structural and functional adaptations increasing insulin release^1,2,3^, beta cell replication^4,5^ and mass^6^ to support energy storage as glycogen and lipids^7,8^. These islet adaptations are highly conserved across species^9,10^. However, prolonged nutrient excess results in beta cell failure, relative insulin insufficiency, and hyperglycemia^11^. Islet architecture^12,13^ and function^14–16^ are strongly modulated by neural inputs. Sympathetic and parasympathetic autonomic nerves innervate islets in multiple species^17^, including humans^18,19^, to coordinate and fine-tune hormone release^15^ and blood flow^20^ in response to nutritional and other stimuli. Despite their biological importance, little is known about the time course and extent of pancreatic innervation adaptation to increased nutrient availability.

Here we identify rapid, circuit-specific and islet-specific adaptations in innervation to short-term increased nutrient availability via mechanisms involving islet Schwann cell-derived neurotrophic factors. We show that high fat diet disrupts islet innervation, contributing to metabolic abnormalities. We further demonstrate that targeted neuromodulation and pharmacological approaches that restore pancreatic neural inputs can reverse metabolic abnormalities associated with nutrient excess.

## Results

### Circuit-specific adaptations in islet innervation with increased nutrient availability

To investigate the time course of nutrient excess on islet innervation architecture, we exploited modified iDISCO+ with immunolabeling for tyrosine hydroxylase (TH), a marker for sympathetic neurons^21^, and vesicular acetylcholine transporter (VAChT), a marker for parasympathetic neurons^22^. Wild-type mice were fed 60% High Fat Diet (HFD) (Fig S1a), a model of nutrient excess that induces obesity (Fig S1c) and metabolic dysfunction (Fig S1 b-o) accompanied by significant changes in islet hormone release (Fig S1p,q) and islet structure (Fig S1r). In mice fed short term, 1-week HFD, we observed a significant increase in the volume of TH+ immunolabeled sympathetic innervation (after excluding TH+ beta cells) of islet glucagon+ volumes and of islet insulin+ volumes in the splenic region of the pancreas consistent with sympathetic hyperinnervation (Fig 1a-h). In contrast, there were no significant changes in parasympathetic VAChT+ innervation of pancreatic alpha or beta cells at this time point (Fig 1i-o). In keeping with previous studies^23,24^, TH+ and VAChT+ exocrine pancreatic innervation density was lower than endocrine pancreas innervation. In addition, we found there were no diet-induced changes in exocrine innervation with short-term high fat diet (Fig S2a,d). With 4 weeks of HFD, the effects of diet on TH+ innervation of islet glucagon+ volumes were no longer significant, but a trend to increased sympathetic innervation of islet insulin+ volumes in the splenic region of the pancreas persisted (Fig 2a-h). There were no effects of 4 weeks HFD on VAChT+ innervation volume of islet insulin+ volumes or islet glucagon+ volumes (Fig 2i-o) and exocrine innervation was not significantly altered by HFD (Fig S2b,e).

**Figure 1.**
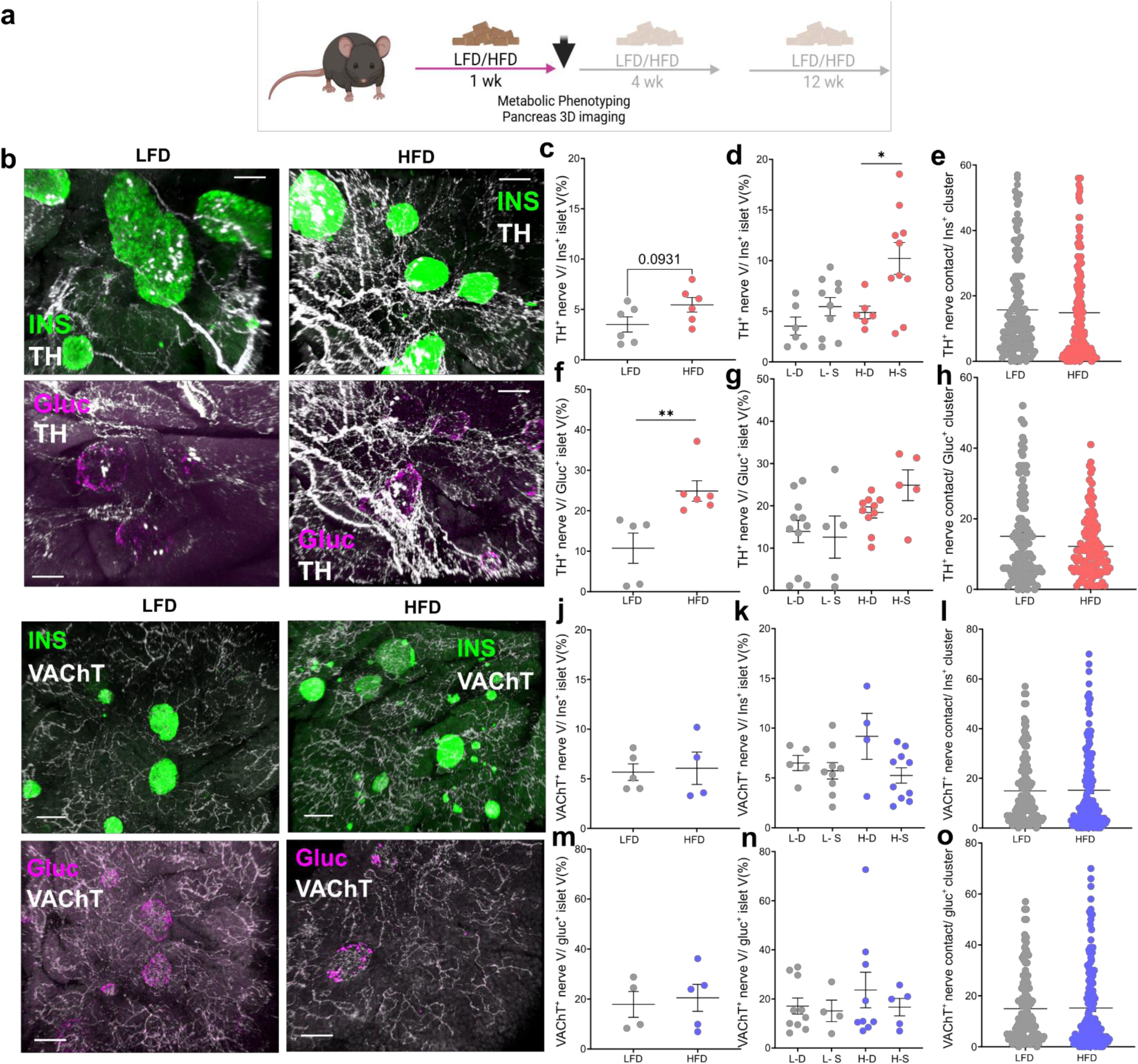
Analysis of islet autonomic innervation after short term HFD. a) Experimental design scheme b) Representative confocal images of iDISCO+ cleared pancreatic sympathetic innervation (TH, white) and insulin (green) or glucagon (magenta), 1 week after HFD exposure (right panels) and LFD diet control (left panels), Scale bars, 100 μm. TH+ (Sympathetic) nerve volume within insulin+ beta cells as a percentage of insulin+ beta cell volume. Each data point represents the average innervation across duodenal and splenic regions of pancreas. LFD: n=6 HFD: n=6. Mann-Whitney test (c) and comparing duodenal (D) vs. splenic (S) regions. L-S: LFD Splenic, L-D: LFD Duodenal, H-S: HFD Splenic, H-D: HFD duodenal. N=6-10/Group. Unpaired t-test, * P<0.05. control duodenal, C-S: control splenic, H-D: HFD duodenal, H-S: HFD splenic. LFD: 6-10 HFD: n= 6-10 (d). e) Contacts made between TH+ nerve surfaces and insulin+ beta cell clusters. LFD: n=6HFD: n=6. f) TH+ (Sympathetic) nerve volume within glucagon+ alpha cells as a percentage of glucagon+ alpha cell volume. Each data point represents the average innervation across duodenal and splenic regions of pancreas. LFD: n=5 HFD: n=6. Mann-Whitney test g) TH+ (Sympathetic) nerve volume within glucagon+ alpha cells as a percentage of glucagon+ alpha cell volume. comparing duodenal (D) vs. splenic (S) regions. N=5-11/Group. h). Contacts made between TH+ nerve surfaces and glucagon+ alpha cell clusters. LFD: n=5 HFD: n=6. i) Representative confocal images of iDISCO+ cleared pancreatic parasympathetic innervation (VAChT, white) and insulin (green) or glucagon (magenta), 1 week after HFD exposure (right panels) and LFD diet control (left panels), Scale bars, 100 μm. j) VAChT+ nerve volume within insulin+ beta cells as a percentage of insulin+ beta cell volume. LFD: n=5 HFD: n=4 k) VAChT+ nerve volume within insulin+ beta cells as a percentage of insulin+ beta cell volume and comparing duodenal (D) vs. splenic (S) regions. N=4-10/Group. l) Contacts made between VAChT+ nerve surfaces and insulin+ beta cell clusters. LFD: n=5 HFD: n=4 m). VAChT+ nerve volume within glucagon+ alpha cells as a percentage of glucagon+ alpha cell volume. LFD: n=4 HFD: n=5 n) VAChT+ nerve volume within glucagon+ alpha cells as a percentage of glucagon+ alpha cell volume comparing duodenal (D) vs. splenic (S) regions. N=4-10/Group. o). Contacts made between VAChT+ nerve surfaces and glucagon+ apha cell clusters. LFD: n=4 HFD: All data represented as mean± SEM

**Figure 2.**
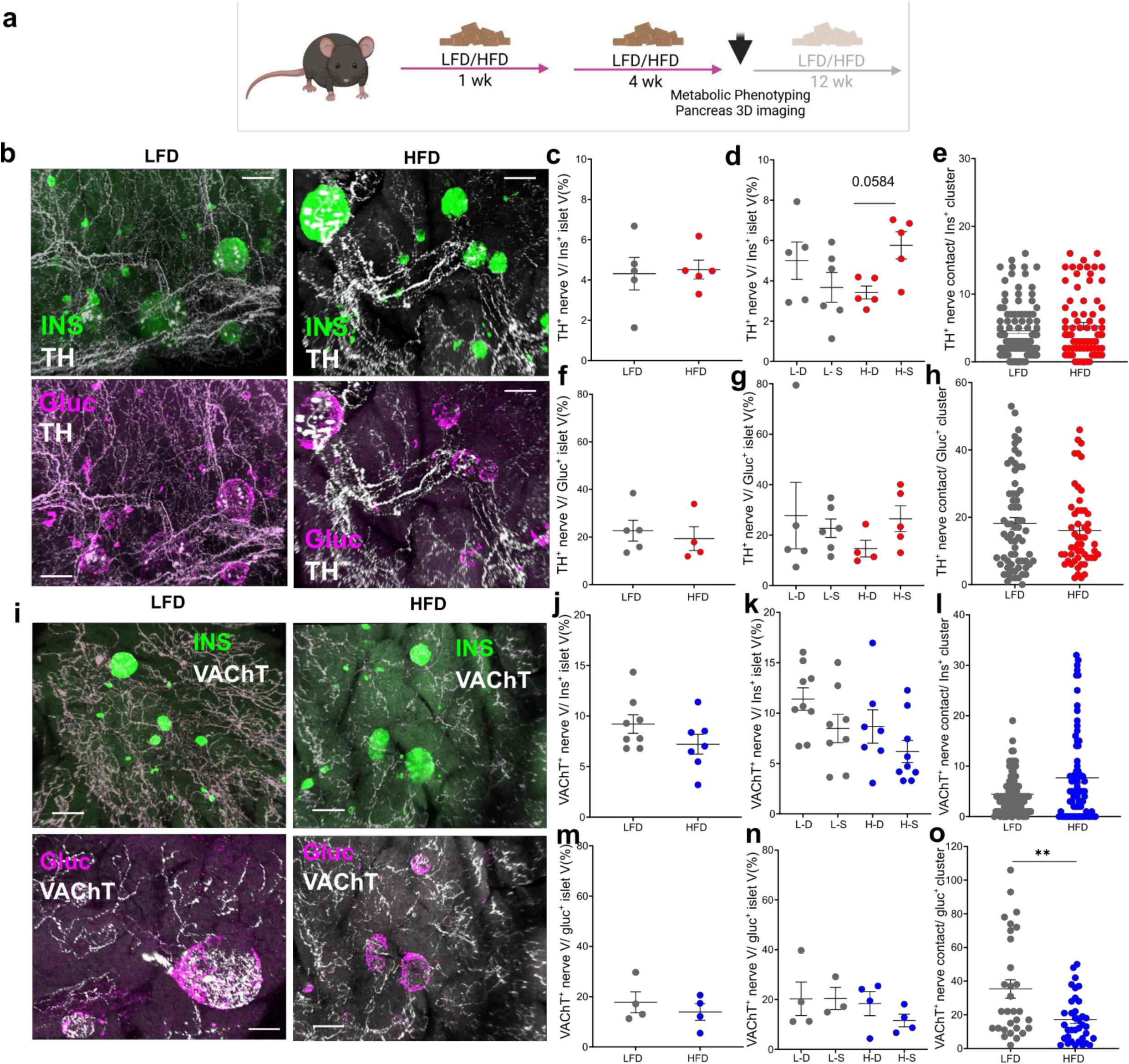
Analysis of islet autonomic innervation after intermediate term HFD. a) Experimental design scheme. b) Representative confocal images of iDISCO+ cleared pancreatic sympathetic innervation (TH, white) and insulin (green) or glucagon (magenta), 1 week after HFD exposure (right panels) and LFD diet control (left panels), Scale bars, 100 μm. TH+ (Sympathetic) nerve volume within insulin+ beta cells as a percentage of insulin+ beta cell volume. Each data point represents the average innervation across duodenal and splenic regions of pancreas. LFD: n=5 HFD: n=5. Mann-Whitney test (c) and comparing duodenal (D) vs. splenic (S) regions. L-S: LFD Splenic, L-D: LFD Duodenal, H-S: HFD Splenic, H-D: HFD duodenal. N=5-6/Group. Unpaired t-test, * P<0.05. control duodenal, C-S: control splenic, H-D: HFD duodenal, H-S: HFD splenic. LFD: 6-10 HFD: n= 6-10 (d). e) Contacts made between TH+ nerve surfaces and insulin+ beta cell clusters. LFD: n=6HFD: n=6. f) TH+ (Sympathetic) nerve volume within glucagon+ alpha cells as a percentage of glucagon+ alpha cell volume. Each data point represents the average innervation across duodenal and splenic regions of pancreas. LFD: n=5 HFD: n=4. Mann-Whitney test g) TH+ (Sympathetic) nerve volume within glucagon+ alpha cells as a percentage of glucagon+ alpha cell volume. and comparing duodenal (D) vs. splenic (S) regions. N=4-6/Group. h). Contacts made between TH+ nerve surfaces and glucagon+ alpha cell clusters. LFD: n=5 HFD: n=4 i) Representative confocal images of iDISCO+ cleared pancreatic parasympathetic innervation (VAChT, white) and insulin (green) or glucagon (magenta), 1 week after HFD exposure (right panels) and LFD diet control (left panels), Scale bars, 100 μm. j) VAChT+ nerve volume within insulin+ beta cells as a percentage of insulin+ beta cell volume. LFD: n=8HFD: n=7. k) VAChT+ nerve volume within insulin+ beta cells as a percentage of insulin+ beta cell volume comparing duodenal (D) vs. splenic (S) regions. N=7-9/Group. l) Contacts made between VAChT+ nerve surfaces and insulin+ beta cell clusters. LFD: n=5 HFD: m) VAChT+ nerve volume within glucagon+ alpha cells as a percentage of glucagon+ alpha cell volume. LFD: n=4 HFD: n=4 n) VAChT+ nerve volume within glucagon+ alpha cells as a percentage of glucagon+ alpha cell volume comparing duodenal (D) vs. splenic (S) regions. N=3-4/Group o) Contacts made between VAChT+ nerve surfaces and glucagon+ apha cell clusters. LFD: n=4HFD: n=4. All data represented as mean± SEM

After 12 weeks of HFD, when mice had marked hyperglycemia and profound metabolic dysfunction (Fig S1), we observed significant circuit-specific and islet-specific alterations in innervation. HFD did not significantly alter TH+ innervation density of alpha or beta cells but significantly increased TH+ contacts with both beta and alpha cell clusters (Fig 3a-h). In contrast, VAChT+ innervation volume in islet insulin+ volumes (normalized to islet volume) was significantly reduced, particularly in the splenic region of the pancreas, with significantly fewer contacts between VAChT+ fibers and both insulin+ beta and alpha cell clusters (Fig 3i-o). In keeping with other time points, prolonged HFD did not significantly alter exocrine pancreas innervation (Fig S2c,f).

**Figure 3.**
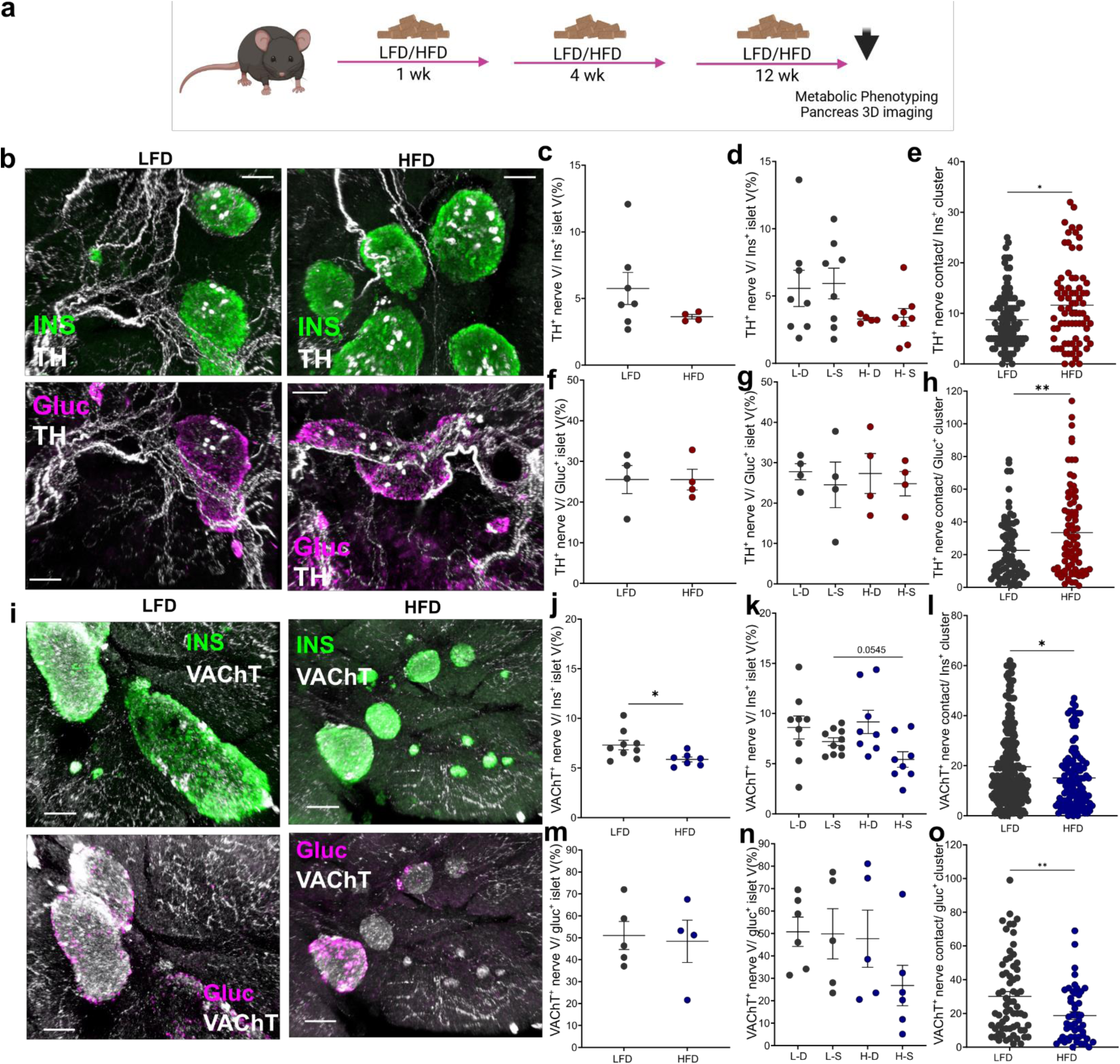
Analysis of islet autonomic innervation after 12 weeks HFD. a) Schema of experimental design. b) Representative confocal images of iDISCO+ cleared pancreatic sympathetic innervation (TH, white) and insulin (green) or glucagon (magenta), after 12-weeks LFD (control, left panels) or HFD (right panels). Scale bars, 100 μm. c) TH+ (Sympathetic) nerve volume within insulin+ beta cells as a percentage of insulin+ beta cell volume. Each data point represents the average innervation across duodenal and splenic regions of pancreas. LFD: n=7 HFD: n=4. d) TH+ (Sympathetic) nerve volume within insulin+ beta cells as a percentage of insulin+ beta cell volume in duodenal (D) vs. splenic (S) pancreatic regions. L-S: LFD Splenic, L-D: LFD Duodenal, H-S: HFD Splenic, H-D: HFD duodenal. N=5-8/Group. e) Contacts made between TH+ nerve surfaces and insulin+ beta cell clusters. LFD: n=7, HFD: n=4. Mann-Whitney test, * P<0.05. f) TH+ (Sympathetic) nerve volume within glucagon+ alpha cells as a percentage of glucagon+ alpha cell volume. Each data point represents the average innervation across duodenal and splenic regions of pancreas. LFD: n=4HFD: n=4. g) TH+ (Sympathetic) nerve volume within glucagon+ alpha cells as a percentage of glucagon+ alpha cell volume in duodenal (D) vs. splenic (S) pancreatic regions. N=4/Group. h) Contacts made between TH+ nerve surfaces and glucagon+ alpha cell clusters. LFD: n=5 HFD: n=4. Mann-Whitney test, ** P<0.01. i) Representative confocal images of iDISCO+ cleared pancreatic parasympathetic innervation (VAChT, white) and insulin (green) or glucagon (magenta), after 12-week LFD (control, left panels) or HFD (right panels). Scale bars, 100 μm. j) VAChT+ nerve volume within insulin+ beta cells as a percentage of insulin+ beta cell volume. LFD: n=9 HFD: n=7. Mann-Whitney test, * P<0.05. k) VAChT+ nerve volume within insulin+ beta cells as a percentage of insulin+ beta cell volume comparing duodenal (D) vs. splenic (S) pancreatic regions. N=8-9/Group. Mann-Whitney test, L-S vs H-S. l) Contacts made between VAChT+ nerve surfaces and insulin+ beta cell clusters. LFD: n=5 HFD: n=4. Mann-Whitney test, * P<0.05. m) VAChT+ nerve volume within glucagon+ alpha cells as a percentage of glucagon+ alpha cell volume. LFD: n=5HFD: n=4. n) VAChT+ nerve volume within glucagon+ alpha cells as a percentage of glucagon+ alpha cell volume comparing duodenal (D) vs. splenic (S) pancreatic regions, N=5-6/Group. O) Contacts made between VAChT+ nerve surfaces and glucagon+ apha cell clusters. LFD: n=4 HFD: n=4. Mann-Whitney test, ** P<0.01. All data represented as mean± SEM.

Together these data reveal a previously unrecognized rapid plasticity of pancreatic innervation, with significantly increased islet sympathetic innervation in response to short-term nutrient excess that progresses to significant islet parasympathetic neuropathy with prolonged metabolic challenge.

### Altered pancreatic innervation contributes to metabolic dysfunction

High fat diet results in adaptive changes in beta cells with increased insulin secretion and beta cell proliferation^1,3–5^. However, this is insufficient to overcome peripheral insulin resistance resulting in relative insulin deficiency and hyperglycemia that becomes more pronounced with prolonged high fat diet^11^. As pancreatic sympathetic and parasympathetic innervation modify islet hormone release and blood glucose^14,15,25,26^, we hypothesized that HFD-induced plasticity in pancreatic innervation may contribute to metabolic dysfunction. We first used chemogenetic neuromodulation to investigate whether increased pancreatic sympathetic drive with short term high fat diet may contribute to metabolic dysfunction. Chemogenetic targeting of pancreas-projecting sympathetic neurons was achieved by co-injecting retrograde-traveling AAV8 expressing cre recombinase into the pancreas and AAVPHP.S-DIO-hSyn-hM3DGq into the celiac ganglia of the same mice (Fig 4a,b). In mice on a control low fat diet (LFD) for 1 or 4 weeks, pancreatic sympathetic activation did not significantly alter basal glucose or glucose tolerance (Fig 4c-h) though plasma glucagon was significantly increased by sympathetic activation during GTT after 1 week of LFD (Fig S3a-d). With HFD, basal glucose and glucose tolerance were not further impaired by DREADD-mediated stimulation of pancreatic sympathetic neurons (Fig 4c-e). Pancreatic sympathetic activation after 4 weeks of HFD did not significantly increase basal glucose or blood glucose levels during GTT but the fold change in glucose with GTT was significantly enhanced by sympathetic activation (Fig 4f-h, Fig S3b). These data suggest that diet-induced increases in sympathetic innervation after 1 week of HFD may occlude any further response to their chemogenetic stimulation while increasing sympathetic activity at 4 weeks HFD elicits subtle impairments in glucose tolerance. In keeping with the reported effects of sympathetic activation in response to hypoglycemia, sympathetic stimulation significantly impaired insulin sensitivity in 1 week LFD-fed mice (Fig 4i, j). However, surprisingly, chemogenetic activation of sympathetic innervation in 1- and 4-week HFD-fed mice significantly lowered blood glucose during insulin tolerance testing compared to mice expressing mCherry in pancreatic sympathetic innervation (Fig 4i-l). Together, these data suggest increased sympathetic drive may contribute to increased glycemic variability with high fat diet, a risk factor for both diabetes and complications of diabetes^27,28^.

**Figure 4.**
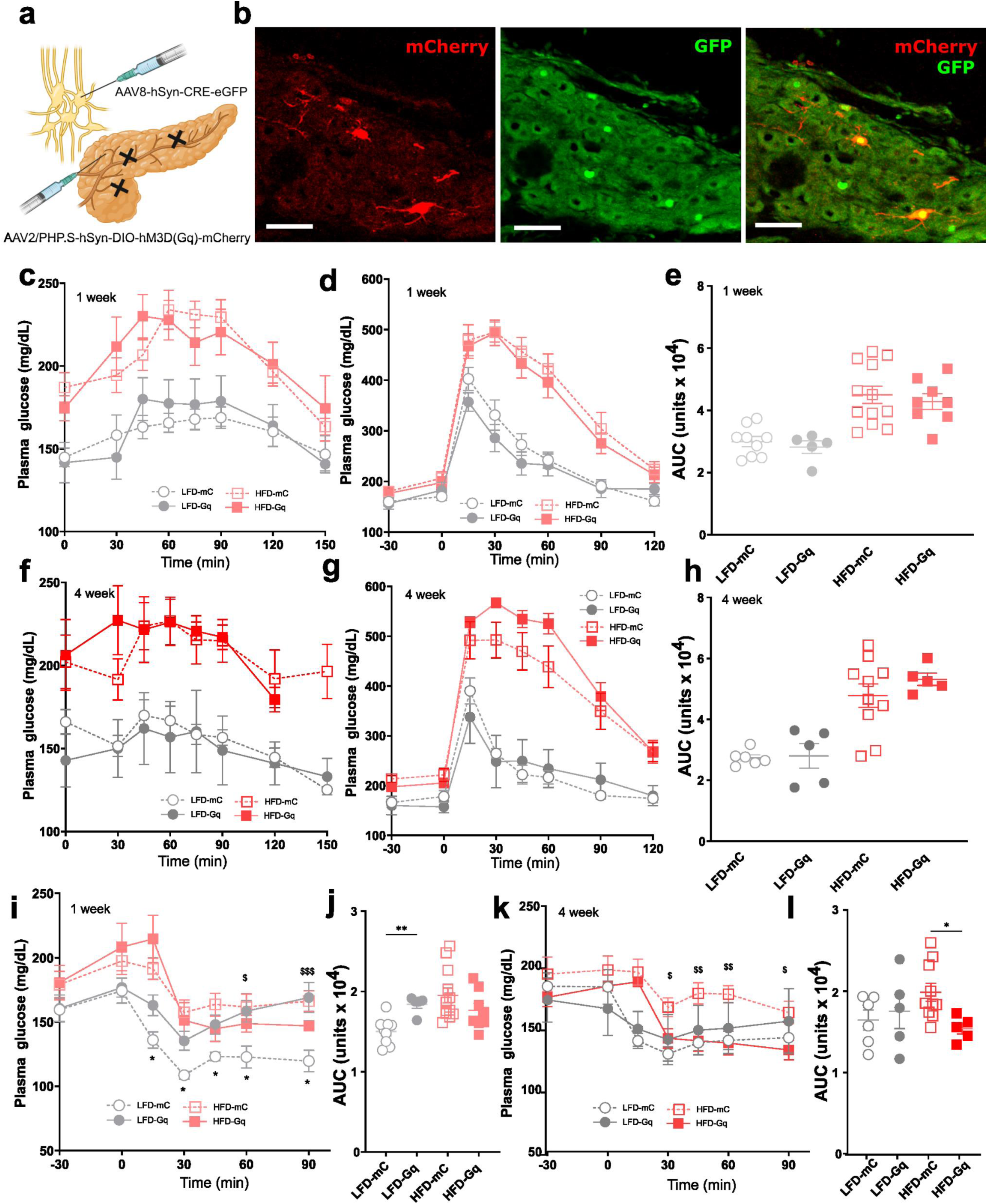
Effects of pancreatic sympathetic activation in glucose homeostasis. a) Schema of viral delivery for pancreatic sympathetic activation b) Images of mCherry (red) and GFP (green) expression in sympathetic pancreas-projecting neurons in the CG after intracoeliac injection of AAV-PHPs-hSyn-DIO-hM3D(Gq)-mCherry and intrapancreatic injection of AAV8-hSyn-CRE-eGFP. Sacel bar 70 µm. Effects of CNO (i.p., 3mg/kg) in AAVdj-hSyn-DIO-hM3D(Gq)-mCherry/AAV8-hSyn-CRE-eGFP mice compared to AAV8-hSyn-DIO-mCherry/AAV8-hSyn-CRE-eGFP mice (control) after 1 week of LFD (LFD-mC N= 9, LFD-Gq N=5) or HFD LFD (HFD-mC N= 12, HFD-Gq N=8) on c) Blood glucose in 6hr fasted mice, d) Glucose Tolerance Test (2g/kg, i.p) and e) Cumulative blood glucose during GTT (AUC, 0 min to 120 min). i)Insulin tolerance test (Insulin 0.3 U/kg i.p.). Mixed-effect analysis * P<0.05 LFD-mC vs LFD-Gq, at 15’, 30’, 45’ 60’, 90’. Mixed-effect analysis $ P<0.05 HFD-mC vs HFD-Gq at 60’, $$$ P< 0.001 at 90’. j) Cumulative blood glucose during ITT (AUC, 0 min to 120 min). Unpaired t-test ** P<0.01 LFD-mC vs LFD-Gq. Effects of CNO (i.p., 3mg/kg) in AAVdj-hSyn-DIO-hM3D(Gq)-mCherry/AAV8-hSyn-CRE-eGFP mice compared to AAV8-hSyn-DIO-mCherry/AAV8-hSyn-CRE-eGFP mice (control) after 4 weeks of LFD or HFD (LFD-mC N= 9, LFD-Gq N=5) or HFD LFD (HFD-mC N= 12, HFD-Gq N=8) on f) Blood glucose in 6hr fasted mice, g) Glucose Tolerance Test (2g/kg, i.p) and h) Cumulative blood glucose during GTT (AUC, 0 min to 120 min). k) Insulin tolerance test (Insulin 0.3 U/kg i.p.). Mixed-Effect analysis HFD-mC vs HFD-Gq, $ P<0.05 at 30’ and 90’, $$ P<0.01 at 45’ and 60’. l) Cumulative blood glucose during ITT (AUC, 0 min to 120 min). Unpaired t-test * P<0.05 HFD-mC vs HFD-Gq. All data represented as mean± SEM

We next investigated the functional consequences of reduced pancreatic parasympathetic innervation that we observed with prolonged high fat diet. To specifically deplete pancreatic parasympathetic innervation, we delivered an adeno-associated virus (AAV) with cre-dependent expression of diphtheria toxin A (AAV-EF1a-mCherry-Flex-DTA) into the pancreatic duct of Chat-IRES-cre mice (Fig 5a). As previous studies suggest decreased pancreatic parasympathetic innervation contributes to cephalic phase insulin release^29,30^, we first examined the effects of pancreatic parasympathetic nerve depletion on blood glucose in fed mice. In mice on LFD, reduced pancreatic parasympathetic innervation significantly increased blood glucose in the fed state at 1,4 and 12 weeks and increased blood glucose in 6 hour fasted mice after 12 weeks of LFD (Fig 5a-f). Further, depletion of pancreatic parasympathetic innervation impaired glucose tolerance in mice after 12 weeks of LFD (Fig 5g-i). By contrast, DTA-mediated parasympathetic nerve depletion did not further alter glycemic regulation in mice on a HFD where parasympathetic innervation is already depleted (12-weeks), or at earlier time points (1-week or 4-week HFD) perhaps in keeping with functional deficits in pancreatic parasympathetic innervation before structural remodeling (Fig 5g-l). Loss of pancreatic parasympathetic innervation did not significantly affect insulin sensitivity or plasma insulin responses to ipGTT (Fig S4).

**Figure 5.**
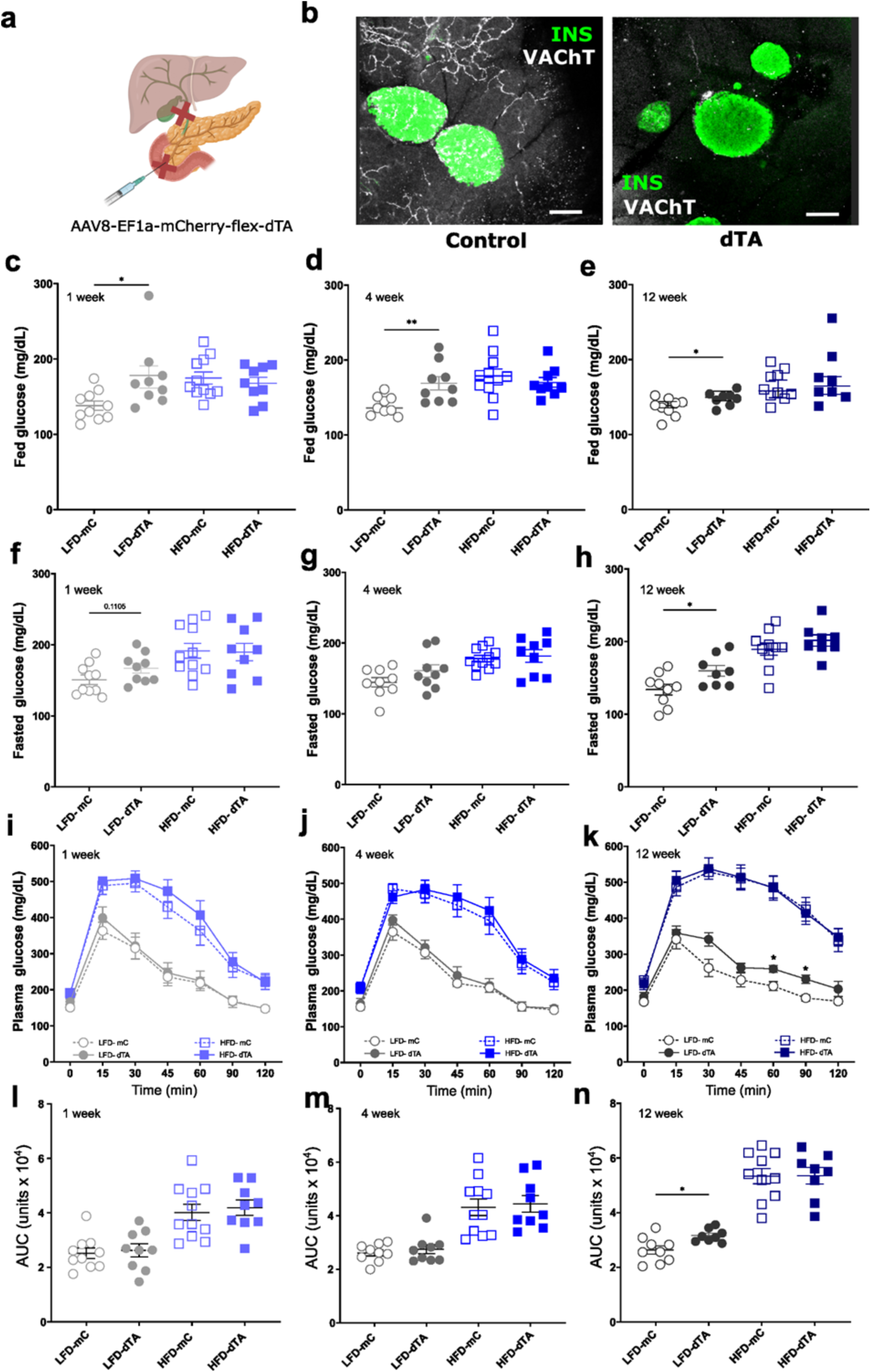
Effects of parasympathetic pancreatic innervation ablation in HFD. a) Schema of viral delivery for pancreatic parasympathetic depletion b) Representative confocal images of iDISCO+ cleared pancreatic parasympathetic innervation (VAChT (white)) and insulin (green) 4 weeks after intraductal delivery of AAVPHP.S-EF1a-mCherry-flex-dtA or AAV8-EF1a-DIO-mCherry in ChAT-IRES-CRE mice. Scale bars, 100 μm. Effects of parasympathetic pancreatic innervation depletion in ChAT-IRES-cre mice expressing AAV8-EF1a-mCherry-flex-dtA or AAV8-EF1a-DIO-mCherry on LFD (LFD-dTA, LFD-mC) or on HFD (HFD-dTA, HFD-mC) on c) Blood glucose levels in fed mice in early light phase after 1 week LFD or HFD. (LFD-mC N=10, LFD-dTA N=9, HFD-mC N=10, HFD-dTA N=9).Two tailed t-test, * P<0.05 LFD-mC vs LFD-Gq d) Blood glucose levels in fed mice in early light phase after 4 weeks LFD or HFD. (LFD-mC N=9, LFD-dTA N=9, HFD-mC N=10, HFD-dTA N=9). Two tailed t-test, ** P<0.01 LFD-mC vs LFD-Gq e) Blood glucose levels in fed mice in early light phase after 12 weeks LFD or HFD.(LFD-mC N=9, LFD-dTA N=8, HFD-mC N=10, HFD-dTA N=8). Two tailed t-test, P<0.05 LFD-mC vs LFD-Gq f) Blood glucose in 6 hour fasted mice after 1 week LFD or HFD. Two tailed t-test. LFD-mC vs LFD-Gq g) Blood glucose in 6 hour fasted mice after 4 weeks LFD or HFD. h) Blood glucose in 6 hour fasted mice after 12 weeks LFD or HFD.Two tailed t-test, * P<0.05 LFD-mC vs LFD-Dta. i) Glucose Tolerance Test (ipGTT, 2g/kg glucose) after 1 week of LFD or HFD (LFD-mC N=10, LFD-dTA N=9, HFD-mC N=10, HFD-dTA N=9). j) Glucose Tolerance Test after 4 weeks of LFD or HFD (LFD-mC N=9, LFD-dTA N=9, HFD-mC N=10, HFD-dTA N=9) k) Glucose Tolerance Test after 12 weeks of LFD or HFD (LFD-mC N=9, LFD-dTA N=8, HFD-mC N=10, HFD-dTA N=8). Two-way repeated measures ANOVA with post-hoc Tukey’s multiple comparison test (* P< 0.05, 60’ and 90’ LFD-mC vs LFD-dTA). l) Cumulative blood glucose during GTT (AUC, 0 min to 120 min) after 1 week of LFD or HFD. m) Cumulative blood glucose during GTT (AUC, 0 min to 120 min) after 4 weeks of LFD or HFD. n) Cumulative blood glucose during GTT (AUC, 0 min to 120 min) after 12 weeks of LFD or HFD. *P<0.05. Two-tailed t-test, LFD-mC vs LFD-dTA. All data represented as mean± SEM

Together, these results indicate that interventions which mimic high fat diet-induced changes in pancreatic innervation - increased sympathetic drive and reduced parasympathetic innervation – likely contribute to impaired glucose homeostasis.

### Pancreatic innervation can ameliorate high fat diet-induced metabolic dysfunction

Given our findings that 1-week exposure to HFD increased islet sympathetic innervation and that enhanced sympathetic activity amplified glycemic variability, we hypothesized that reducing pancreatic sympathetic inputs in mice with short-term nutrient excess would improve glucose homeostasis. Using pancreas-targeted delivery of the catecholamine-specific neurotoxin, 6 hydroxydopamine^31^ (6-OHDA), we specifically depleted intrapancreatic TH+ fibers, but not TH+ beta cells, in adult mice on short-term, 1 week LFD and HFD (Fig 6a,b). While plasma glucose was not significantly different in fed mice, depletion of pancreatic sympathetic innervation significantly lowered blood glucose in 6 hour fasted, HFD-fed mice and improved glucose tolerance compared to vehicle-treated mice fed HFD for 1 week (Fig 6c-f), without significant differences in insulin sensitivity (Fig 6g,h). In mice fed LFD, pancreatic sympathetic nerve depletion did not alter fed blood glucose, 6 hr fasted blood glucose, or glucose tolerance (Fig 4c-f). However, blood glucose levels following an insulin challenge were significantly lower in mice with loss of pancreatic TH+ fibers compared to vehicle-treated mice (Fig 6g,h), in keeping with known effects of pancreatic sympathetic innervation in the counter-regulatory response to hypoglycemia^32^. These results suggest that reducing sympathetic innervation in early HFD significantly improves glycemic control.

**Figure 6.**
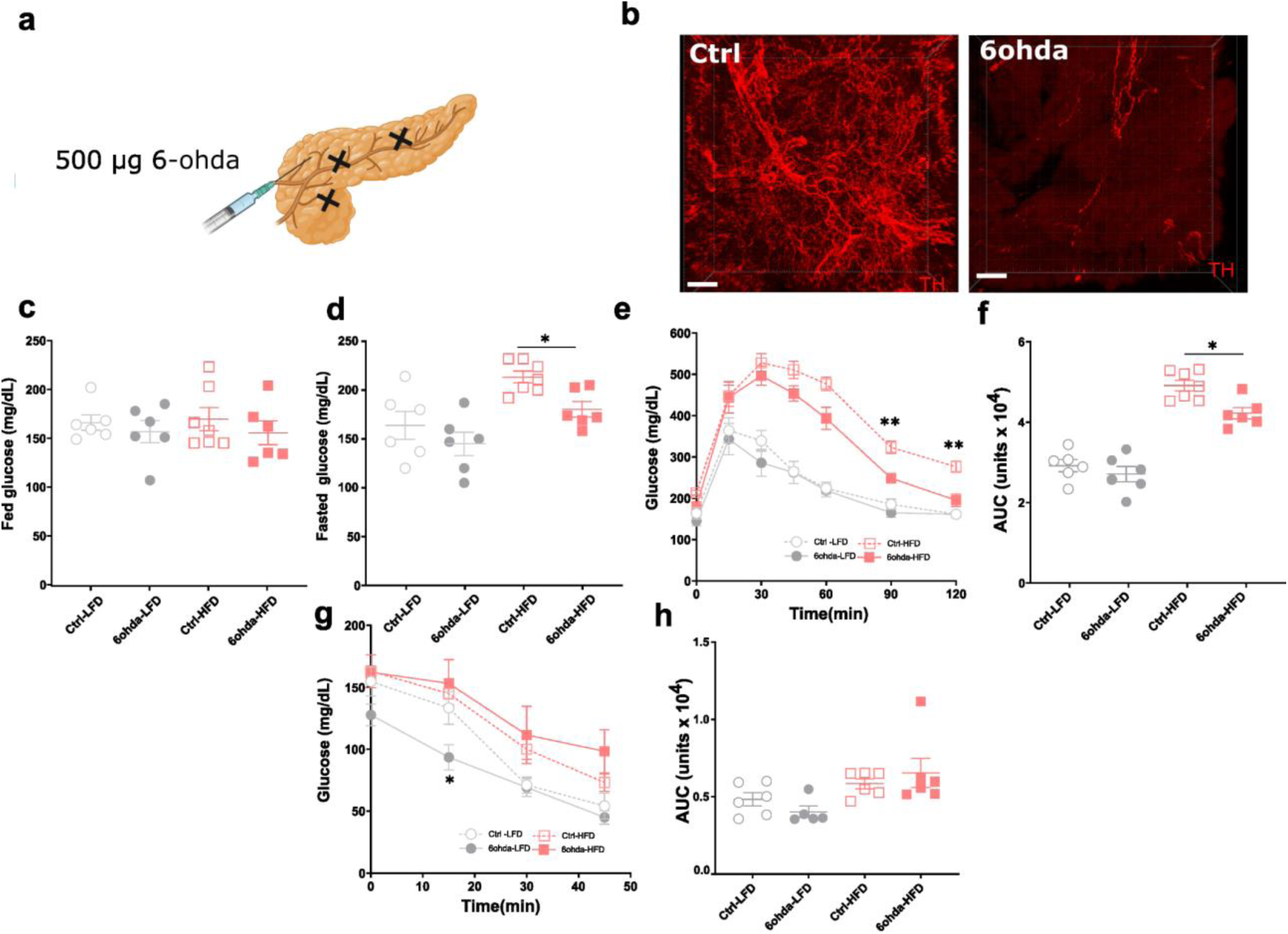
Effects of sympathetic pancreatic innervation ablation in HFD. a) Schema of 6 hydroxydopamine (6-ohda) delivery for pancreatic sympathetic depletion b) Representative confocal images of iDISCO+ cleared pancreatic sympathetic innervation (TH, red). Sace bar 100 μm. Effects of pancreatic sympathetic innervation depletion after 1 week LFD or HFD (6ohda-LFD N= 6, 6ohda-HFD N= 6) compared to vehicle-treated control mice (Ctrl-LFD N= 7, Ctrl-HFD N= 6) on c) Blood glucose levels in fed mice in early light phase d) Blood glucose in 6 hour fasted mice. Brown-Forsythe and Welch ANOVA tests with Dunnett’s T3 multiple comparison test, * P <0.05 Ctrl-HFD vs 6ohda-HFD. e) Glucose Tolerance Test (ipGTT, 2g/kg glucose) 2-way Anova with Tukey’s multiple comparison test, ** P<0.01, Ctrl-HFD vs 6ohda-HFD at 90’ and 120’. f) Cumulative blood glucose during GTT,(AUC, 0-120 min). 2-way Anova with Tukey’s multiple comparison test, *P,0.05 Ctrl-HFD vs 6ohda-HFD. g) Insulin Tolerance Test (0.75 u/kg, ip) Mixed-effect analysis * P<0.05 Ctrl-LFD vs 6ohda-LFD at 15’. h) Cumulative blood glucose during ITT (AUC 0-45) All data represented as mean± SEM

Previous studies have reported that chemogenetic and optogenetic activation of pancreatic parasympathetic innervation can improve insulin release and glycemic control in wild-type and insulin deficient mice with normal insulin sensitivity^15,25,26^. Therefore, we investigated whether activating pancreatic cholinergic nerve activity was sufficient to improve metabolic responses in insulin-resistant mice after short and long-term nutrient excess. Chemogenetic activation was achieved using intraductal delivery of AAV expressing cre-dependent hM3Gq in Chat-IRES-cre mice (Fig 7a,b). Pancreatic parasympathetic nerve activation lowered blood glucose in 6 hour fasted mice after 1 and 4 weeks of LFD, and improved glucose tolerance in insulin-sensitive, LFD-fed mice at all time points without resulting in hypoglycemia (Fig 7c, d, f, g, i, j, l,m). After 1 and 4 weeks of HFD treatment, when parasympathetic innervation is not significantly reduced, pancreatic parasympathetic nerve activation also normalized blood glucose in 6 hr fasted mice and improved glucose tolerance (Fig 7c, d, f, g, i, j, l, m). Unexpectedly, even in mice with parasympathetic neuropathy after 12 weeks of HFD, parasympathetic activation restored basal blood glucose and glucose tolerance to levels seen in LFD control mice (Fig 7e,h,k,n). In LFD-fed mice, parasympathetic activation significantly increased plasma insulin compared to baseline in response to a glucose challenge (Fig 8a-c). However, parasympathetic stimulation also significantly increased plasma glucagon, most markedly at 1 week (Fig 8d-f). In HFD-fed mice, parasympathetic nerve stimulation significantly increased plasma insulin compared to mCherry-expressing control mice at 1 week and significantly increased plasma insulin compared to baseline in response to a glucose challenge at 4 and 12 weeks (Fig 8a-c), without significant effects on plasma glucagon (Fig 8d-f).

**Figure 7.**
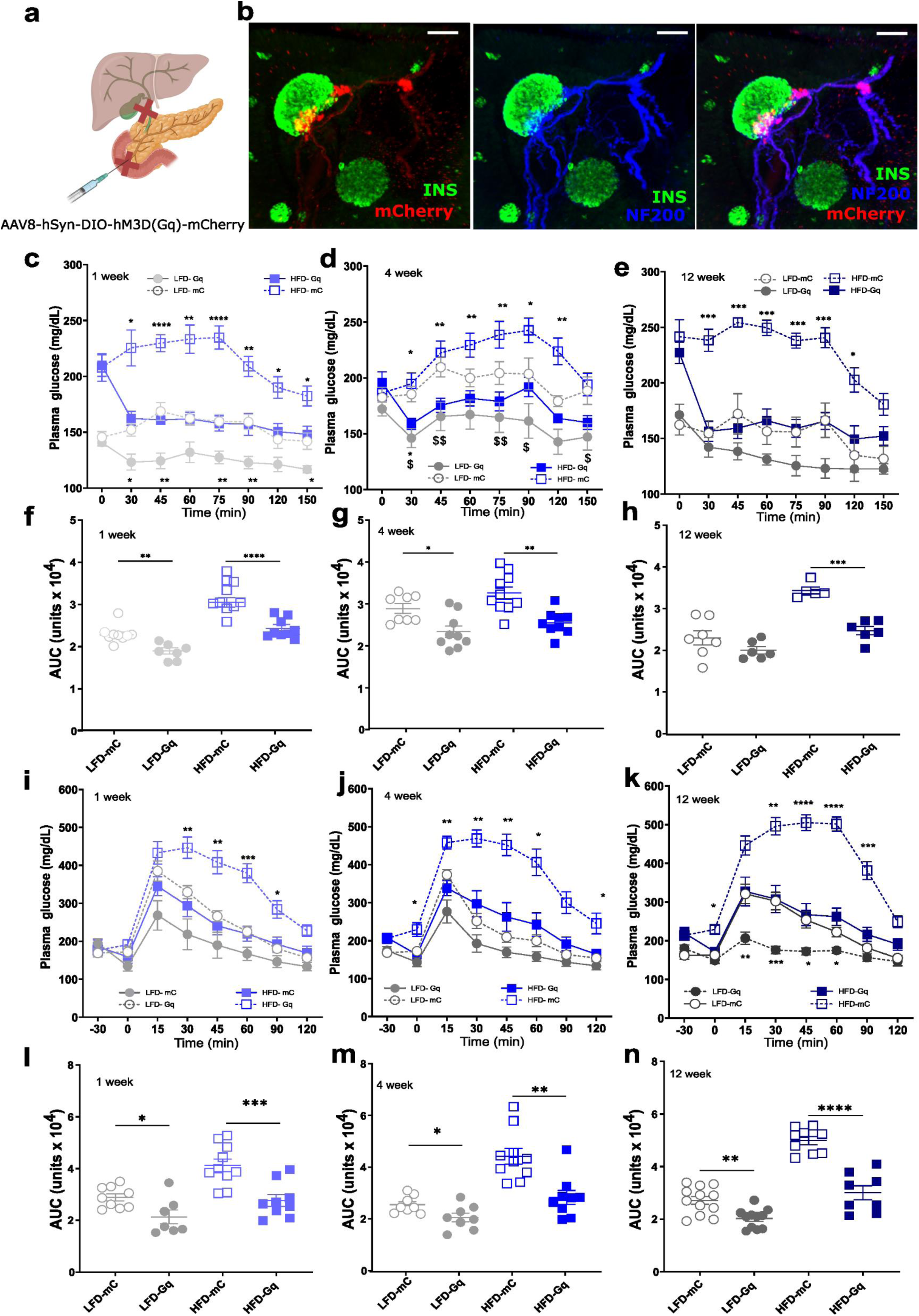
Pancreatic parasympathetic activation improves glucose control in HFD. a) Schema of viral delivery for pancreatic parasympathetic innervation activation b) Representative confocal images of iDISCO+ cleared pancreatic tissue with mCherry-tagged viral expression (mCherry, red), parasympathetic innervation (VAChT (blue)) and insulin (green) after intraductal delivery of AAV8-hSyn—DIO-hM3D(Gq)-mCherry in ChAT-IRES-CRE mice. Scale bars, 100 μm. Effects of CNO treatment (3mg/kg, i.p.) ChAT-IRES-cre mice expressing AAV8-hSyn-DIO-hM3D(Gq)-mCherry compared to AAV8-EF1a-DIO-mCherry exposed to LFD (LFD-Gq, LFD-mC) or HFD (HFD-Gq, HFD-mC) on c) Blood glucose levels in 6 hr fasted mice after 1 week LFD or HFD (N= 7-10/group). Two-way ANOVA, with post-hoc Tukey’s multiple comparison test * P<0.05, **P<0.01, *** P<0.001, **** P<0.0001 d) Blood glucose levels in 6 hr fasted mice after 4 weeks LFD or HFD (N= 8-10/group). Two-way repeated measures ANOVA with post-hoc Tukey’s multiple comparison test, * P<0.05, **P<0.01, *** P<0.001, **** P<0.0001. Two-way repeated measures ANOVA with post-hoc Tukey’s multiple comparison test $ P<0.05, $$ P<0.01 LFD-Gq vs LFD-mC e) Blood glucose levels in 6 hr fasted mice after 12 weeks LFD or HFD (N= 7-10/group). Two-way ANOVA with post-hoc Tukey’s multiple comparison test, * P<0.05, **P<0.01, *** P<0.001, **** P<0.0001 f) Cumulative blood glucose during basal glucose testing after 1 week LFD or HFD. One way ANOVA with post-hoc Tukey’s multiple comparison test, * P<0.05, **P<0.01, *** P<0.001, **** P<0.0001 g) Cumulative blood glucose during basal glucose testing after 4 weeks LFD or HFD. One way ANOVA post-hoc Tukey’s multiple comparison test, * P<0.05, **P<0.01, *** P<0.001, **** P<0.0001 h) Cumulative blood glucose during basal glucose testing after 12 weeks LFD or HFD. One way ANOVA post-hoc Tukey’s multiple comparison test, * P<0.05, **P<0.01, *** P<0.001, **** P<0.0001 i) Glucose Tolerance Test (ipGTT, 2g/kg glucose) after 1 week of LFD or HFD. Two-way ANOVA with Bonferroni’s multiple comparison test, * P<0.05, **P<0.01, *** P<0.001, **** P<0.0001 HFD-Gq vs HFD-mC j) Glucose Tolerance Test (ipGTT, 2g/kg glucose) after 4 weeks of LFD or HFD. Two-way ANOVA, with post-hoc Tukey’s multiple comparison test, * P<0.05, **P<0.01, *** P<0.001, **** P<0.0001 HFD-Gq vs HFD-mC k) Glucose Tolerance Test (ipGTT, 2g/kg glucose) after 12 weeks of LFD or HFD. Two-way ANOVA with post-hoc Tukey’s multiple comparison test,, * P<0.05, **P<0.01, *** P<0.001, **** P<0.0001 l) Cumulative blood glucose during GTT (AUC, 0 min to 120 min) after 1 week of LFD or HFD. Two-tailed t-test *P<0.05 LFDmC vs LFD-Gq, ***P<0.001 HFD-mC vs HFD-Gq m) Cumulative blood glucose during GTT (AUC, 0 min to 120 min) after 4 weeks of LFD or HFD. Two-tailed t-test *P<0.05 LFD-mC vs LFD-Gq, **P<0.01 HFD-mC vs HFD-Gq n) Cumulative blood glucose during GTT (AUC, 0 min to 120 min) after 12 weeks of LFD or HFD. *P<0.05. Two-tailed t-testt-test **P<0.01 LFD-mC vs LFD-Gq, ****P<0.0001 HFD-mC vs HFD-Gq All data represented as mean± SEM

**Figure 8.**
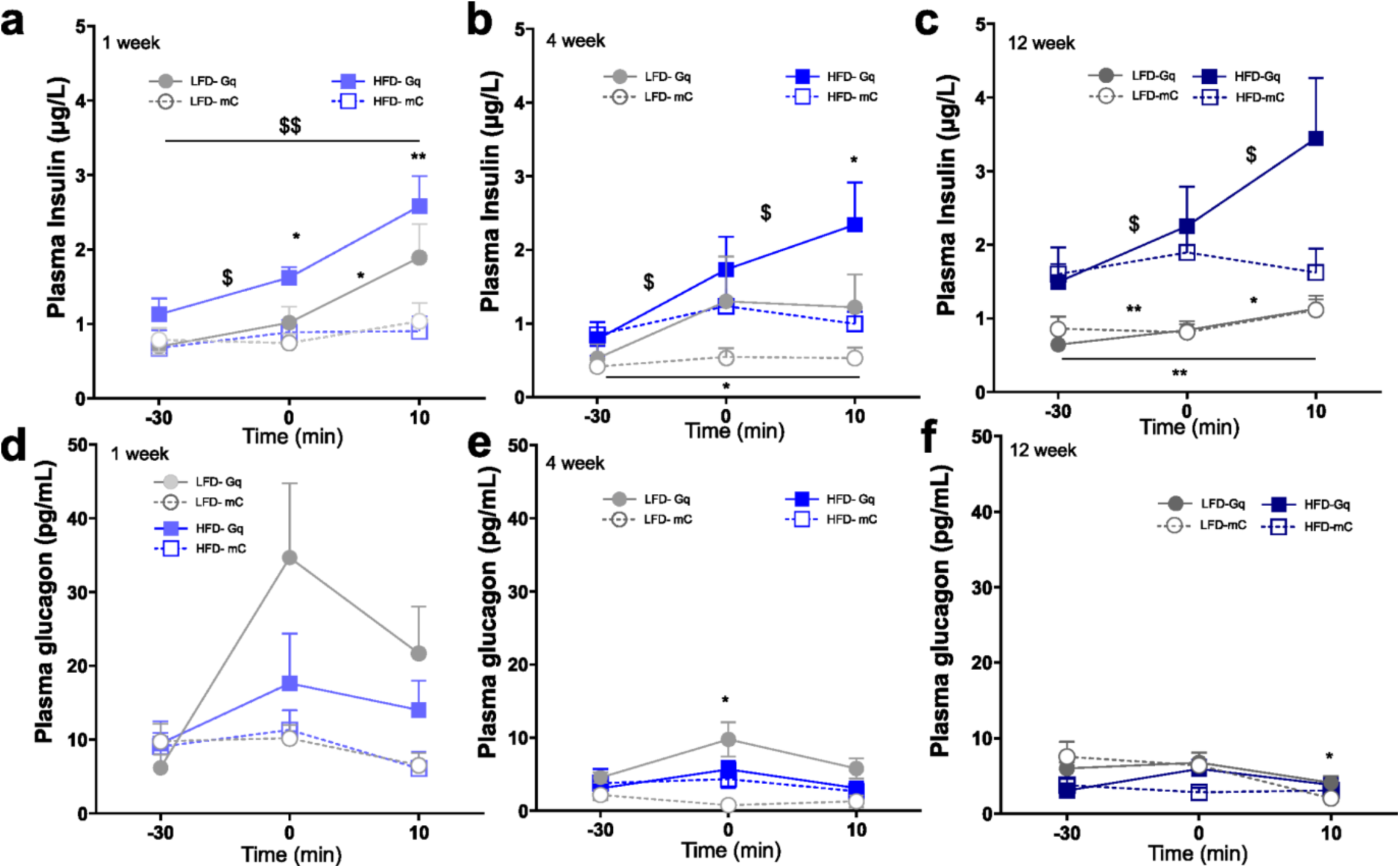
Effects of Pancreatic parasympathetic activation on hormones regulating glucose control in HFD. Effects of CNO treatment (3mg/kg, i.p.) ChAT-IRES-cre mice expressing AAV8-hSyn-DIO-hM3D(Gq)-mCherry compared to AAV8-EF1a-DIO-mCherry on LFD (LFD-Gq, LFD-mC) or on HFD (HFD-Gq, HFD-mC) on a) Plasma insulin levels during GTT in at −30, 0 and 10 min after 1 week of LFD or HFD. (N=7-9 group). Two-way ANOVA, with post-hoc Sidak’s multiple comparison test, * P<0.05, **P<0.01, HFD-Gq vs HFD-mC. Two-way ANOVA, with post-hoc Tukey’s multiple comparison test $ P<0.05 HFD-Gq −30’ to 0, * P<0.05 LFD-Gq 0’ to 10’. b) Plasma insulin levels during GTT in at −30, 0 and 10 min after 4 weeks of LFD or HFD. (N=8-10/group). Two-way ANOVA, with post-hoc Bonferroni’s multiple comparison test, * P<0.05 HFD-Gq vs HFD-mC. Two-way ANOVA, with post-hoc Tukey’s multiple comparison test $ P<0.05 HFD-Gq −30’ to 0, $ P<0.05 HFD-Gq −0’ to 10’, * P<0.05 LFD-Gq −30’ to 10’. c) Plasma insulin levels during GTT in at −30, 0 and 10 min after 12 weeks of LFD or HFD. (N=7-12 group). Mixed-effect analysis, with post-hoc Tukey’s multiple comparison test $ P<0.05 HFD-Gq −30’ to 0’, HFD-Gq 0’ to 10’ ** P<0.01 LFD-Gq - 30’ to 0’, LFD-Gq −30’ to 10’. * P<0.05 LFD-Gq 0’ to 10’. d) Plasma glucagon levels during GTT in at −30, 0 and 10 min after 1 week of LFD or HFD. (N=7-10 group). e) Plasma glucagon levels during GTT in at −30, 0 and 10 min after 4 weeks of LFD or HFD. (N=8 group). Mixed-effect analysis with post-hoc Tukey’s multiple comparison test, * P<0.05 LFD-Gq vs LFD-mC. f) Plasma glucagon levels during GTT in at −30, 0 and 10 min after 12 weeks of LFD or HFD. (N=7-12 group). All data represented as mean± SEM

These findings suggest that short-term increases in pancreatic parasympathetic activity are sufficient to enhance glucose-stimulated insulin release without hypoglycemia, a feature which could improve the safety profile of glucose-lowering therapies^33^. Published studies suggest the actions of parasympathetic stimulation are mediated by the M3 muscarinic receptor^34^, however the effects of hyperglycemia and high fat diet on muscarinic receptor expression are conflicting^35,36^. Therefore, we first examined cholinergic and adrenergic receptor expression in murine pancreatic alpha and beta cells after HFD treatment for 16 weeks in a published data set^37^. Only cholinergic M3 receptor, nicotinic beta 2 receptor and alpha-2A adrenergic receptor RNA were detectable in beta and alpha cells, with significant upregulation of M3 receptors on beta cells from mice with HFD (Fig 9a). We next examined whether cevimeline, an FDA-approved M3/M1 receptor agonist used for the treatment of Sjorgren’s syndrome^38^, might also be effective in improving glucose homeostasis in obese, hyperglycemic mice. cevimeline (10mg/kg, i.p.) significantly improved glucose tolerance and increased plasma insulin in obese, hyperglycemic mice on HFD (Fig 9b-e). Similarly, oral administration of cevimeline (10mg/kg by oral gavage) significantly improved glucose tolerance in obeses HFD-fed mice (Fig 9f,g). These data suggest that M3/M1 receptor agonist treatment, already approved for use in humans, may be effective as an antidiabetic agent.

**Figure 9.**
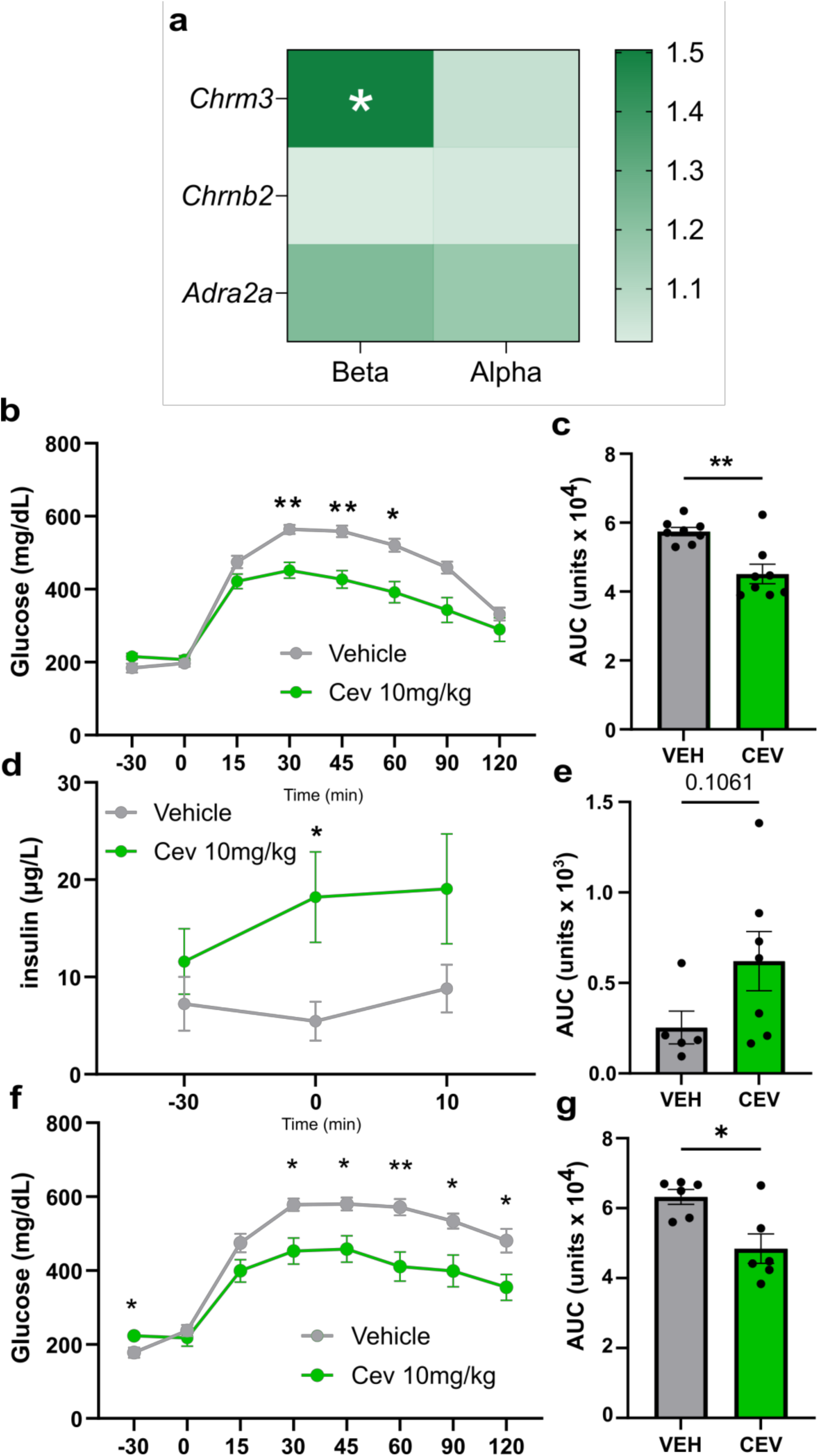
M1/M3 agonist improves glucose control in HFD. a) Heatmap of fold change between LFD and 16 weeks HFD in cholinergic and adrenergic gene expression in alpha and beta cells in reanalyzed data from^37^ b) Intraperitoneal Glucose Tolerance Test (ipGTT, 2g/kg glucose) in cevimeline-treated mice (i.p., 10 mg kg^−1^), after 6 hr fast on mice exposure to 16 weeks of HFD (N=8/ group). Two-way ANOVA, with post-hoc Sidak’s multiple comparison test, **P<0.01, *P<0.05 CEV vs VEH. c) Cumulative blood glucose during GTT (Area under the Curve, 0-120 min). Man-Whitney test, ** P<0.01 *P<0.05 CEV vs VEH. d) Plasma Insulin levels during GTT in at −30, 0 and 10 min in Cevimeline treated mice (N=5-7/group). Mixed-effect analysis with post-hoc Tukey’s multiple comparison test, * P<0.05 CEV vs VEH. e) Cumulative plasma insulin during GTT (Area under the Curve, −30 to 10 min). Man-Whitney test f) Intraperitoneal Glucose Tolerance Test (ipGTT, 2g/kg glucose) in mice treated with Cevimeline orally (i.p., 10 mg kg^−1^), after 6 hr fast on mice exposure to 16 weeks of HFD (N=8/ group). Two-way ANOVA, with post-hoc Sidak’s multiple comparison test, **P<0.01, *P<0.05 CEV vs VEH. g) Cumulative blood glucose during GTT (Area under the Curve, 0-120 min) in f). Man-Whitney test, *P<0.05 CEV vs VEH. All data represented as mean± SEM

### Schwann cell S100b regulates pancreatic hyperinnervation in early metabolic challenge

As HFD results in rapid circuit-specific and islet-specific remodeling of innervation, we assessed whether alterations in local neuroguidance cues might contribute to early increases in sympathetic innervation of islets. To address this, we first reanalyzed our published bulk RNA sequencing data^39^ from islets isolated from C57Bl6 mice treated with normal chow or 45% high fat diet for 1 week (Fig 10a). Using the gene cards database, we identified 4971 genes related to “neurite extension” and 2835 genes related to “neurotrophic factor” of which 4448 and 2299 genes respectively were identified as expressed in the RNAseq data set from pancreatic islets. Filtering for significantly upregulated genes (FC>Log2 1.5 and p < 0.05), we identified 15 unique genes that were upregulated by short-term high fat diet (Fig 10 b,c). For unbiased assessment of which islet cell types might be expressing these neuroguidance cues, we used PanglaoDB^40^, a publicly available database of cell type markers. Our search, using all significantly upregulated neurotrophic/neurite extension genes, revealed the major cell types expressing these genes were neurons and glia, including Schwann cells, a glial cell type present in the peripheral nervous system and around pancreatic islets^41^ (Fig 10d).

**Figure 10.**
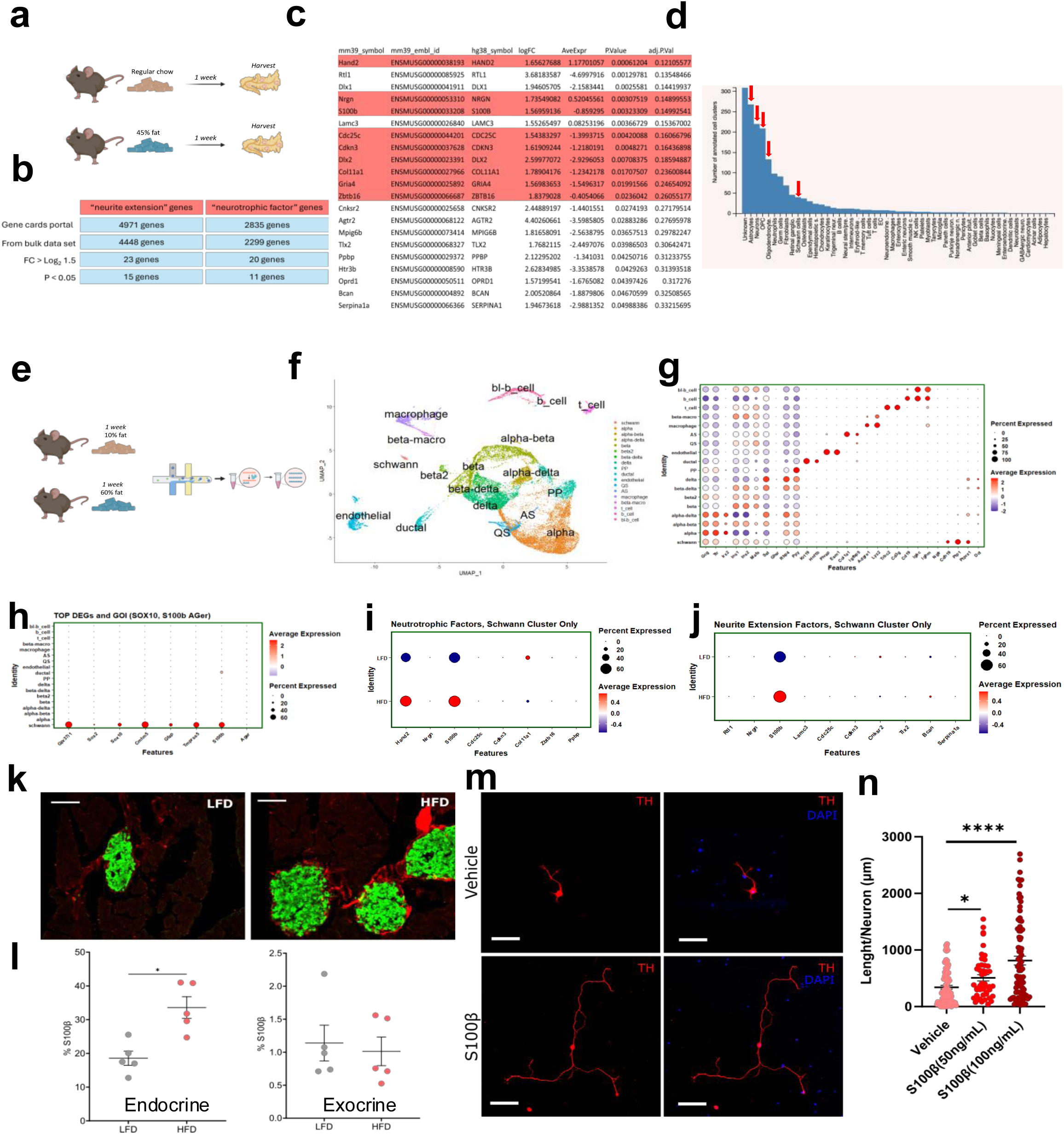
Neurotrophic factors from pancreatic islet Schwann cells induce sympathetic nerve growth. a) Schema of bulk RNAseq studies from islets harvested from mice after 1 week of normal chow or 45% HFD. b) Table of the numbers of neurotrophic and neurite extension genes from the Gene cards portal, expressed in bulk RNAseq data, with a fold change (FC) greater than 1.5 fold in HFD group, and with P value < 0.05. c) Table of neurotrophic factor and neurite extension genes upregulated in islets from HFD-fed mice. Highlighted genes are present in lists of neurotrophic factors and neurite extension factors. d) Cell type annotation of upregulated genes using PanglaoDB e) Schema of single cell RNAseq studies from mice after 1 week of LFD or 60% HFD from reference 41 f) UMAP plot of all islet cells identified by scRNAseq, showing the distribution of clusters and cell types. g) Expression of genes in each cluster used for annotation h) Top differentially expressed genes in Schwann cell cluster i) Expression of neurotrophic factor genes in Schwann cell cluster from islets harvested from mice after 1 week of LFD or HFD j) Expression of neurite extension genes in Schwann cell cluster from islets harvested from mice after 1 week of LFD or HFD k) Immunohistochemistry for S100b protein (red) and insulin (green) in pancreatic tissue from mice after 1 week of LFD or HFD. Scale bar 100 μm. l) Quantification of S100b area in peri-islet region and exocrine pancreas (N= 5/group). * P<0.05, Mann-Whitney test m) Representative immunocytochemistry images for tyrosine hydroxylase (red) and DAPI (blue) in cultured primary sympathetic neurons from coeliac ganglia cultured with vehicle or S100b Scale bar 100 μm. n) Effects of 24-hour incubation with S100b 50ng/ml (2 independent replicates) or 100ng/ml (3 independen replicates) on neurite length per primary sympathetic neuron. One way ANOVA with post-hoc Tukey’s multiple comparison test, *P<0.05, ****P<0.0001. All data represented as mean± SEM

To determine whether HFD modulates the expression of neuroguidance cues in Schwann cells, we reanalyzed a published single cell RNA sequencing data set from C57Bl6 mice fed LFD or 60% HFD for 1 week^42^, conditions in line with those used in our studies (Fig 10e). Reannotation identified 18 distinct cell types, including islet endocrine cell types, macrophages, and immune cells as well as an “undefined” population. Further analysis of this population revealed significant expression of Schwann cell markers: *Ngfr*, *Cdh19*, *Plp1*, *Ptprz1*, and *Dct.* To confirm this, we examined the top differentially expressed genes in this cluster and found enriched expression of *Gpr37l1, Sox2, Sox10, Cmtm5, Gfap, Tmprss5* and *S100b*, canonical Schwann cell markers (Fig 10 f-h). Next, we assessed expression of the neurotrophic/neurite extension genes identified in our bulk RNAseq data in the scRNAseq data set with short-term LFD or HFD treatment. Expression of both *Hand2,* a transcription factor implicated in maintaining sympathetic innervation, and the calcium binding protein, *S100b* were increased in pancreatic Schwann cells from mice fed short-term HFD (Fig 10 i-j). To validate these findings, we examined S100b protein in pancreatic tissue using immunohistochemistry. In keeping with increased gene expression, HFD significantly increased islet S100b immune-positive area but not exocrine S100b+ area suggesting specific changes in S100b protein in islet Schwann cells in response to short-term HFD diet (Fig 10 k,l) but not long-term HFD (Fig S5b). To probe the effects of S100b on sympathetic neurons from the celiac ganglia which innervate the pancreas, we examined neurite length in cultured mouse primary sympathetic neurons in response to S100b treatment. Short-term (24 hr) treatment with S100b dose-dependently increased primary sympathetic neuron neurite length (Fig 10m, n).

These findings suggest that HFD induces S100b expression in peri-islet Schwann cells resulting in increased islet sympathetic innervation that contributes to increased glycemic variation and metabolic dysfunction.

### Summary

Metabolic challenges such as high fat diet rapidly alter pancreatic islet function and structure, but longitudinal data to determine its impact on the detailed architecture and function of specific autonomic circuits innervating the islet are largely unknown. Neural circuits play important roles in the regulation and coordination of islet function^14–16^ with feeding, hypoglycemia and stress. However, prior studies using 2D imaging or examining catecholamine turnover in response to HFD have been conflicting with reports of increased^2^, decreased^43^, or unchanged^44^ pancreatic innervation/tone in genetically obese or HFD-fed rodents. Our findings identify circuit-specific plasticity in pancreatic autonomic innervation that partly contributes to the glycemic abnormalities that are characteristic of HFD-fed mice. Surprisingly, the structure of islet autonomic innervation was remodeled without effects on exocrine innervation. Our findings provide compelling evidence that the highly localized plasticity is, in part, through upregulation of the neurotrophic factor S100b in pancreatic Schwann cells that encapsulate islets. Unexpectedly, counteracting the HFD-induced functional changes in islet innervation using either neuromodulation or FDA-approved cholinergic agonists, improved glycemic control in both lean, normoglycemic and obese, hyperglycemic mice without inducing hypoglycemia. If these findings extend to humans, they suggest that targeting dysregulated islet autonomic circuits, for example with repurposed FDA-approved therapies, may provide an additional pathway to improve metabolic dysfunction in obesity.

## MATERIALS AND METHODS

### Viral Plasmids and Constructs

The following adeno-associated viruses were purchased commercially from Addgene (Watertown, MA): AAV8-hSyn-DIO-hM3D(Gq)-mCherry (44361-AAV8), AAV8-hSyn-DIO-mCherry (50459-AAV8).

AAV8-EF1a-mCherry-flex-dtA, AAV8-EF1a-DIO-mCherry, AAV2/PHP.S-hSyn-DIO-hM3D(Gq)-mCherry and AAV8-hSyn-CRE-eGFP were obtained from “Canadian Neurophotonics Platform Viral Vector Core Facility (RRID:SCR_016477, construct-387, construct-kd2, construct-246, and construct-890 respectively)

#### In Vitro Studies

Primary culture of Coeliac Ganglia (CG) sympathetic neurons was adapted from^45^. Briefly, postnatal day 3 pups from WT C57BL/6J mice (Jackson Laboratories, Bar Harbor ME; #000664) were sacrificed and CG was isolated and placed in dissection media Leibovitz L-15 (Thermo Fisher Scientific, 11415064) supplemented with 1% Penicillin-Streptomycin (Gibco; 15140-122) and 8% FAF-BSA (Calbiochem, 126609). Then, CG was incubated with Hanks’ Balanced Salt Solution (HBBS) (Thermo Fisher Scientific, 14185) supplemented with 1mg/mL Collagenase type 2 (Worthington Biochemical, 4176) and 5mg/mL of Dispase II (Roche, 4942078001) for 45 minutes at 37c. Digested CG were triturated by gentle pipetting and placed in Poly-D-Lysine (Sigma-Aldrich P0899) treated chamber slides (Thermo Fisher, 154453). The culture media consisted of: DMEM (Low Glucose, Thermo Fisher Scientific 11885) /Ham’s F12(Thermo Fisher Scientific, 11765) (1:1) supplemented with 5% FAF-BSA, 0.5 mM L-Glutamine (company; G7513), 1% Penicillin-Streptomycin, Insulin-Selenium-Transferrin (100X) (Thermo Fisher Scientific, 41400-045), and 0.125 ug/mL of Nerve Growth Factor (NGF, 1156-NG/CF, R&D). 24 hours after plating, CG were treated with Cytosine-β-D-arabinofuranoside (Sigma-Aldrich, C1768) at 2uM to prevent growth of non-neuronal cells. Finally, S100b (Millipore, 559290) treatment was performed at 50 ng/mL and 100ng/mL for 24h.

Afterwards, cells were fixated using 10% formalin (Fisher Scientific, SF98-4) for 10 min at RT. Fixed cells were stained for Tyrosine Hydroxylase (TH, Millipore, AB152, dilution 1:500, 4c, overnight) and subsequent secondary antibody used was Alexa Fluor 647 anti-rabbit (Jackson ImmunoResearch, 711-605-152). Slides were mounted using Fluoromount mounting media containing DAPI (Southern Biotech, Birmingham, AL; 0100-20). Fixed cells were imaged using an inverted Zeiss LSM 880 confocal microscope at a magnification of 20X. Exposure times and fluorescence intensity were maintained between samples. To measure neurite length, Image analysis was performed using the FIJI plugin NeuroJ.

#### Animal Studies

WT C57BL/6J (Jackson Laboratories; #000664) and heterozygous ChAT-IRES-Cre (Jackson Laboratories; #028861) aged 8-12 weeks were maintained on a 12h/12h light-dark cycle with controlled humidity and temperature, with access to food *ad libitum*. All animal studies were performed with the approval of, and in accordance with, guidelines established by the Icahn School of Medicine at Mount Sinai, and principles of laboratory animal care were followed.

##### High fat diet

Mice were fed a high fat diet (HFD, Research Diets, D12492, 60% fat) or control diet (LFD, Research Diets D12450J, 10% fat). Body weight and blood glucose were measured for 1 week, 4 weeks, and 12 weeks.

##### Surgical Procedures

For all surgical procedures, mice were initially anesthetized with 3.5% vaporized isoflurane, with percentage decreased to 1.5-2.5% during the procedures. Mice were kept on a heating pad for the duration of the procedure. Animals were given buprenorphine (0.05 mg/kg, IP) every 12h for 3 days following surgery.

Intrapancreatic injections were performed as described by Fassanella^46^; AAVs or other reagents were injected in 1-2µL increments into random areas of the pancreas of WT C57BL/6J male mice.

For intra-coeliac infusions, 2 μl of the correspondent AAV were administered into the CG at 0.2 μl min−1, using a a Nanofil 36G bevelled needle (NF36BV-2, World Precision Instruments) and Silflex tubing (SILFLEX-2, World Precision Instruments) attached to a Nanofil 10 μl syringe (NANOFIL, World Precision Instruments).

Lastly, for intraductal infusions, mice were kept on a liquid diet (Ensure, Abbott laboratories, Chicago IL) 1-3 days prior to surgery. Intraductal infusions were performed as described^47^; briefly, 150 µL of PBS containing the desired AAV titer were infused into the pancreas via cannulation through the duodenum into the hepatopancreatic ampulla and common bile duct, at a rate of 6µLmin-1.

###### Activation of Pancreas-projecting sympathetic efferent nerves

WT C57BL/6J male mice received intra-celiac infusions of AAVPHP.S-hSyn-DIO-hM3D(Gq)-mCherry or AAV8-hSyn-DIO-mCherry (2×10^10^ vg/mouse) and intrapancreatic injections of AAV8-hSyn-CRE-eGFP (5×10^11^ vg/mouse, up to 20µL injected, in 1µL increments into random areas of the pancreas).

###### Depletion of Pancreas-projecting sympathetic efferent nerves

WT C57BL/6J male mice received 500 µg of 6-Hydroxydopamine 6-Hydroxydopamine (6-OHDA) hydrobromide (HelloBio, HB1889) via intrapancreatic injections. 6-OHDA was freshly prepared in 0.01% Ascorbic Acid (Sigma-Aldrich, A92902) and injected immediately.

###### Activation of pancreatic parasympathetic efferent nerves

ChAT-IRES-Cre male mice received intraductal infusions of with AAV8-hSyn-DIO-hM3D(Gq)-mCherry (5×10^11^ vg/mouse) or AAV8-hSyn-DIO-mCherry (5×10^11^ vg/mouse)

###### Depletion of pancreatic parasympathetic efferent nerves

ChAT-IRES-Cre male mice received intraductal infusions of AAV-EF1a-mCherry-flex-dtA (5×10^11^ vg/mouse) or AAV8-EF1a-DIO-mCherry (5×10^11^ vg/mouse)

### Metabolic Studies

#### Timing of surgeries and metabolic testing

For 1week HFD studies, mice that underwent surgeries for AAVs expression were rested for 3 weeks and diet consumption began 4 weeks after surgeries. Hence, 1 week LFD and HFD metabolic testing occurred 5 weeks post-surgery. The same animals were used for 4-week metabolic testing which occurred 8 weeks post-surgery when previous studies demonstrate viral expression is not significantly altered^15^. For 12 week LFD and HFD studies, mice were randomized to LFD or HFD groups and diet consumption began. Mice underwent surgery 7 weeks later. Metabolic studies were performed 5 weeks post-surgery and 12 weeks after the start of diets.

Animals receiving 6-OHDA for sympathetic ablation, were started on HFD 5 days post-surgery and metabolic tests commenced 1 week after.

Blood glucose was measured on tail snips using the Contour glucose meter (Bayer, Leverkusen; GER; 7097C). Baseline measurements and intraperitoneal glucose tolerance tests (IPGTT) were administered in the afternoon after a 6h fast. For chemogenetic modulation studies, clozapine-N-oxide (Hello Bio) 3mg/kg in saline/10% DMSO or vehicle (saline/10% DMSO) was administered at time −30min. For baseline measurements, blood glucose was measured at times −30, 0, 15, and then 30 minutes intervals to 120 min or 240 min. For IPGTT, 2g/kg of D-glucose (Sigma; G8270) in saline was injected at time 0, with blood glucose measured at times −30, 0, 15, 30, 45, 60, 90 and 120min. Plasma insulin and glucagon were measured on blood collected during the IPGTT studies at Time −30 (immediately preceding CNO injection), time 0 (immediately preceding glucose injection), and at 10 min. Insulin tolerance testing (ITT) was performed by injecting insulin (HumulinR, Lilly) at 0.3 U/kg in 6h fasting mice. then glucose measurements were taken at −30, 15, 30, 45, 60, 90 and 120min. ITT in 6-OHDA pancreatic ablation studies was performed using 0.75 U/kg of insulin.

For the cevimeline studies, mice were subjected to HFD for 16 weeks. After 6 hours fast, Cevimeline hydrochloride (Hello Bio, HB1490) was injected at 10 mg/kg 30 min before ipGTT. The same animals were given vehicle the following day for a cross-over study. Plasma insulin was measured on blood collected during the IPGTT studies at Time −30 (immediately preceding Cevimeline injection), time 0 (immediately preceding glucose injection), and at 10 min.

##### Body Composition

EchoMRI (EchoMRI-500 Body composition analyzer) was used to determine whole body composition of live animals. A baseline reading was determined using an empty animal holder tube. Animals were then individually placed in the animal holder tube, and EchoMRI was performed to determine body composition (total fat and lean mass). EchoMRI took place after 1, 4 and 12 weeks of LFD and HFD consumption. Weekly body weights were assessed on all animals.

##### Assays

For plasma insulin and glucagon content, bloods were collected during ipGTT at −30, 0 and 10 minutes using 300uL blood collection tubes for plasma (Sarstedt, 16.444.100). Plasma levels of insulin and glucagon were determined by ELISA (Insulin: Mercodia, Uppsala, SWE; 10-1247-01, Glucagon: 10-1271-01). Plasma glucagon levels are in line with published data using ELISA^48,49^

##### Tissue collection

Depending on the study, the following tissues were collected: CG and pancreas. Tissues were maintained in 4% paraformaldehyde overnight before processing.

###### Tissue processing for cryosectioning

To confirm chemogenetic expression in the intracoeliac infusion studies, CG were immersed in 30% sucrose (Sigma-Aldrich, 50389) in PBS overnight, then embedded in O.C.T compound (ThermoFisher, 23-730-572), frozen at −80 °C and sectioned at 25 μm thickness. Tissues were stained overnight with mCherry (Abcam, ab205402) + tyrosine hydroxylase (TH, Millipore, AB152) at 1:1,000 dilution. Subsequent secondary antibodies used were Alexa Fluor 647 anti-rabbit (Jackson ImmunoResearch, 711-605-152) and Alexa Fluor 594 anti-chicken (Jackson ImmunoResearch, 703-585-155). Tissues were stained with DAPI and coverslipped. Samples were visualized using aan inverted Zeiss LSM 880 confocal microscope at a magnification of 20X.

For the S100β staining studies, pancreas from animals receiving 1 week or 12 weeks of HFD or LFD were processed as described above. Then, sections were stained overnight for S100b (Abcam, ab52642) + Insulin at 1:500 insulin (R&D systems MAB1417). Subsequent secondary antibodies used were Alexa Fluor 647 anti-rabbit (Jackson ImmunoResearch, 711-605-152) and Alexa Fluor 488 anti-rat (ThermoFisher # A-21208).

###### Optical clearing of mouse pancreata

Whole-organ staining and clearing were performed on 1 week LFD/HFD pancreatic samples according to a modified iDisco protocol^23^:

Dissected pancreata were dehydrated [20, 40, 60, 80, and 100% methanol at room temperature (RT)], delipidated [100% dichloro-methane (DCM; Sigma-Aldrich, St. Louis, MO, USA)], and bleached in 5% H2O2 (overnight, 4°C). Pancreata were rehydrated (80, 60, 40, and 20% methanol) and permeabilized [5% dimethyl sulfoxide/0.3 M glycine/0.1% Triton X-100/0.05% Tween-20/0.0002% heparin/0.02% NaN3 in PBS (PTxwH)] for 1 day. Pancreata were then placed in blocking buffer [PTxwH + 3% normal donkey serum (Jackson ImmunoResearch, West Grove, PA, USA)] at 37°C overnight. Samples were incubated with primary antibodies (Supplementary Table 1) in blocking buffer for 3 or 6 days (small pancreatic pieces and hemipancreata, respectively) at 37°C. After five washes with PTxwH at RT (final wash overnight), samples were incubated with secondary antibodies in blocking buffer (1:500) for 3 or 6 days. Samples were washed with PTxwH (five times, RT) and PBS (five times, RT), dehydrated with a methanol gradient, then washed in 100% methanol (three times, 30 min each) and DCM (three times, 30 min each), and then transferred to dibenzyl ether (DBE; Sigma-Aldrich) for the final clearing step.

Whole-organ staining and clearing were performed 4 and 12 week LFD/HFD samples according to Adipoclear^50^: Dissected pancreata were dehydrated 20, 40, 60, 80% MeOH/B1N buffer (Glycine (0.3M), Triton X-100 (0.1% v/v), Sodium azide (0.1% (w/v))for 1 hour each at 4°C and 100% methanol for 1 hour at 4°C. Samples were delipidated with 100% dichloromethane (DCM; Sigma-Aldrich, St. Louis, MO, USA)], then washed with 100% MeoH 1 hour each at 4°C. Samples were then bleached in 5% H2O2 in methanol overnight, 4°C. Pancreata were rehydrated with 80, 60, 40, and 20% methanol/B1N buffer 1 hour each at 4°C. Samples were washed in 100% B1N buffer 2×1 hour. Samples were washed in PTxwH buffer (PBS1x, 0.1% Triton X-100, 0.05% Tween 20, Heparin (2µg/mL)) for 2 hours before proceeding to incubation steps. Samples were incubated with primary antibodies (supplementary table 1) in PTxwH buffer for 5 days (small pancreatic pieces and hemipancreata, respectively) at 37°C. Samples were washed with PTxwH at RT (5 min, 10 min, 15 min, 20 min, 1 h, 2 h, 4 h and overnight), samples were incubated with secondary antibodies in blocking buffer (1:500) min for 5 days. Samples were washed with PtXWH at RT (5 min, 10 min, 15 min, 20 min, 1 h, 2 h, 4 h and overnight)and PBS (5, min, 10 min, 30 min), dehydrated with a 25%, 50%, 75% methanol/H_2_0 gradient for 1 hour each at RT and 100% methanol (three times, 30 min each) and DCM (three times, 1 hour each, or until samples sink to the bottom of the tubes), and then transferred to dibenzyl ether shaking at RT overnight (DBE; Sigma-Aldrich) to clear. DBE was replaced before proceeding to imaging.

**Supplementary table 1.**

| Primary | Dilution | Secondary | Dilution |
| --- | --- | --- | --- |
| Insulin, SIGMA, G2624 | 1:1000 | Alexa Fluor 488 anti-rat | 1:500 (ThermoFisher # A-21208) |
| Glucagon R&D systems MAB1417 | 1:2000 | Alexa Fluor 647 anti-mouse | 1:500 (ThermoFisher A32787) |
| VACHT Synaptic Systems #139103 | 1:500 | Alexa Fluor 594 anti-rabbit | 1:500 (ThermoFisher A21207) |
| TH, Millipore, AB152 | 1:500 | Alexa Fluor 594 anti-rabbit | 1:500 (ThermoFisher A21207) |
| S100B Abcam, ab52642 | 1:500 | Alexa Fluor 594 anti-rabbit | 1:500 (ThermoFisher A21207) |

##### Imaging processing and analysis

Z-stacked optical sections were acquired with an UltraMicroscope II (LaVision BioTec, Bielefeld, Germany; 4x magnification with dynamic focus with using maximum projection filter). Mouse hemi-pancreatic samples were imaged at 4x with dynamic focus and with multiple Z-stacks acquired at 4x with 20% overlap and tiled using the plugin TeraStitcher through the ImSpector Pro software (LaVision BioTec).

Small mouse pancreatic sections were imaged in glass-bottom eight-well chambers (Ibidi, Gräfelfing, Germany) filled with immersion media DBE or ECi and imaged using an inverted Zeiss LSM 880 confocal microscope with a 10× [numerical aperture (NA), 0.3] objective and a step size of 5 mm. Spatial resolution for confocal images acquired at 10× was 1.67 mm by 1.67 mm by 5 mm. 3 z-stack images were taken per hemi pancreas, for duodenal and splenic portions. Volumetric data across the 3 z stack images summed and the resulting volumetric data is represented as one data point for regional distribution figures (i.e., splenic vs. duodenal figures). For graphs indicating volumes across the entire pancreas, data were compiled to represent volumetric data from 3 z stack images from the spleen and 3 z stack images from the duodenal pancreas, and the summation of this volumetric data is represented as one data point per animal.

Imaris (Bitplane AG, Zürich, Switzerland) was used to create digital surfaces covering the islets and innervation (10x images) to automatically determine volumes and intensity data. Volume reconstructions were performed using the surface function with local contrast background subtraction. For detection of islets, the threshold factor corresponded to the largest islet diameter in each sample. For detection of nerves, the threshold factor was set to 12.2 mm. A smoothing factor of 10 mm was used for islets, and a factor of 3.25 mm was used for analysis of islets and nerves. For detection of TH^+^ nerves, TH^+^ b cells were manually removed from the final TH^+^ nerve surface by excluding volumes below 120 mm^3^ residing within insulin^+^ islets and overlapping with insulin staining. The Imaris Distance Transform Matlab XTension function was used to calculate the distance of each islet surface from the innervation surface. Distances of islets are reported as the intensity minimum of the distance transformation channel (intensity 0 = islet touching nerve) for each islet surface to the nerve surface as calculated by the distance transformation operation. In confocal images, digital surfaces were created to cover nerves and individual beta cells, alpha cells, or ganglia. For detection of ganglia, a region of interest was manually created around each individual ganglion to create a digital surface specifically covering cell bodies, but not nerve fibers. The Imaris Distance Transform Matlab XTension was then used as above to determine the distance between ganglia and insulin^+^ islets or the distance between nerves and individual a or b cells with a distance of 0 indicating a nerve contact. Limitations to our analyses of endocrine cell contacts include the following: We may not have captured a/b cells with lower staining intensity; in some cases, we could not separate adjacent endocrine cells and therefore counted multiple adjacent cells as a single cell; our method quantifies the number of endocrine cells contacting nerves but does not allow for quantification of number of contacts per endocrine cell

### Single Cell Sequencing Data acquirement and Processing

We downloaded all single cell RNA sequencing data in AR. Piñeros et al’s study^42^, from their corresponding repository GSE162512 (GSM4953223 to GSM4953229) and processed the FASTQ files into H5 count format with CellRanger V6.1.1 on 10X Cloud platform with mm10-2020 reference genome, and we didn’t include intronic counts for the this pipeline.

### Quality Control and Integration

The potential ambient mRNA contamination was adjusted with SoupX (with 10% contamination estimation). Gene count level quality control was performed by filtering out the genes : less than 500 gene count; less than 250 gene varieties (features); less than 0.8 log10 genes per UMI; and greater than 20% mitochondrial gene ratio which is labeled with “mt-“ prefix. Potential doublets were also algorithmically removed with DoubletFinder package, and we applied conventional 8% doublet formation ratio per cell recovered for this study. Subsequently, 7 single cell data were integrated using Seurat’s SCTrasnform function and we applied default method parameter, and cell identity was defined with conventional cell type markers.

#### Statistical Analysis

Data are expressed as means+/- SEM. Analysis were performed using GraphPad Prism. D’Agostino & Pearson test was used to assess normality. For comparison between two groups, and if data were normally distributed, then statistical difference was assessed using an Unpaired two-tailed t-test. If data were not normally distributed, then a Mann Whitney test was performed to assess differences between two groups.

## Acknowledgements

JREC is supported in part by NIH grant T32AG049688. KD was supported in part by the American Heart Association (grant 18PRE33960254). DE is supported in part by the American Heart Association (25PRE1373198). MJG is supported in part by a pilot and feasibility award from the NIDDK-supported Einstein-Sinai Diabetes Research Center (DRC) (P-30 DK020541). AA was supported by a senior postdoctoral fellowship from the Charles H. Revson Foundation (Grant No. 18-25) and a postdoctoral scholarship from Swedish Society for Medical Research (SSMF). RFH was supported by NIH grant F31DK129016. This work was supported in part by grants from the National Institutes of Health (R01DK124461to SAS) and Department of Defense (W81XWH-20-1-0345). Microscopy and image analysis were performed at the Microscopy core at the Icahn School of Medicine at Mount Sinai. The authors wish to thank the NIDDK supported Einstein-Sinai Diabetes Research Center (DRC) (P-30 DK020541).

Figures were created with BioRender.com

