## Supplementary figures S1-S5 for "Nutrient excess remodels islet autonomic innervation via pancreatic schwann cells"

### Figure S1

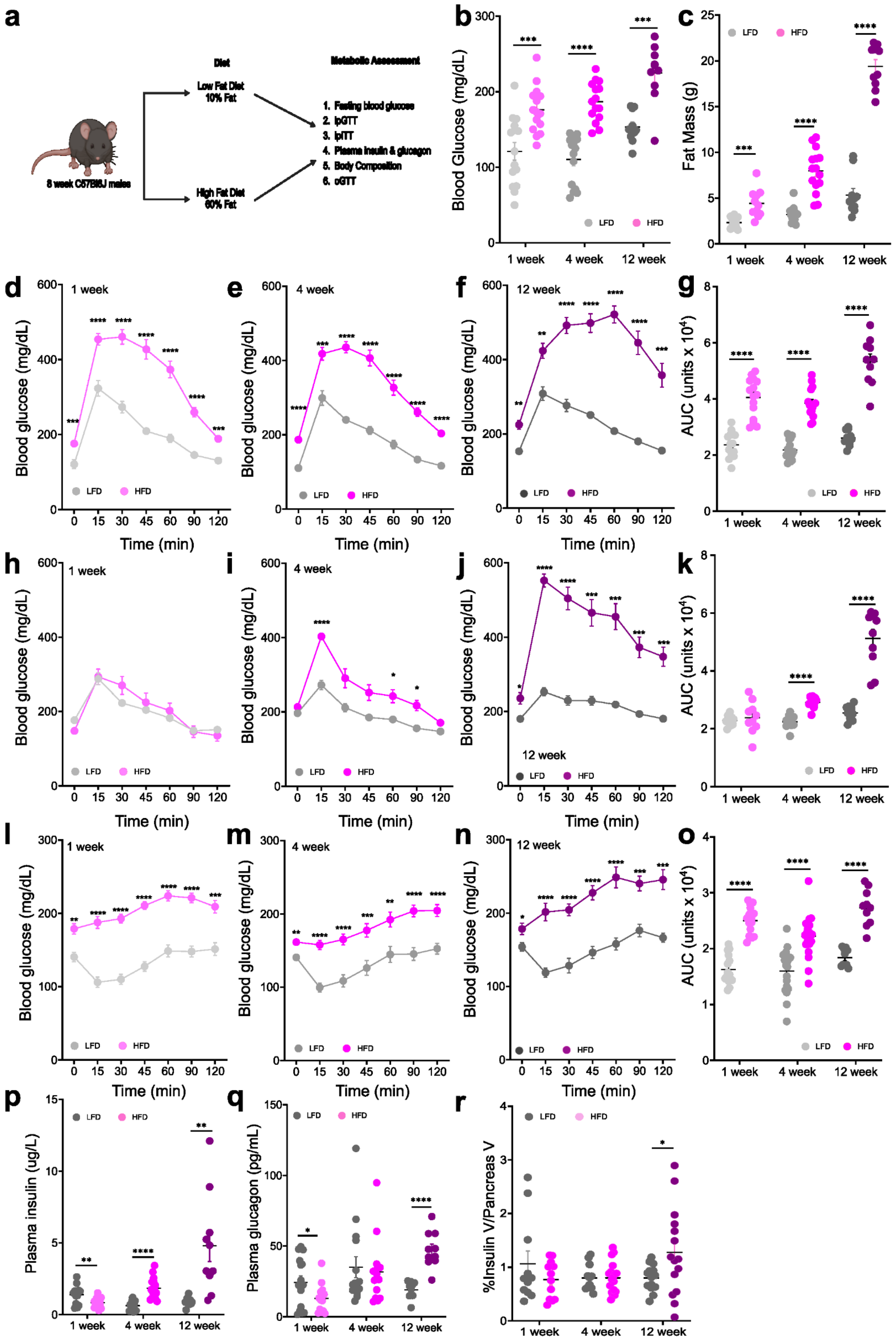

a
Diet
Low Fat Diet 10% Fat
High Fat Diet 60% Fat
8 week C57Bl6J males
Metabolic Assessment
1. Fasting blood glucose
2. ipGTT
3. ipITT
4. Plasma insulin & glucagon
5. Body Composition
6. oGTT
b
Blood Glucose (mg/dL)
1 week
4 week
12 week
LFD
HFD
c
Fat Mass (g)
d
1 week
Blood glucose (mg/dL)
Time (min)
e
4 week
f
12 week
g
AUC (units x 10^4)
h
i
j
k
l
m
n
o
p
Plasma insulin (ug/L)
q
Plasma glucagon (pg/mL)
r
%Insulin V/Pancreas V

**Figure S1. Effects of HFD consumption on glucose metabolism**
a) Experimental design scheme.
b) Blood glucose levels (6 hours fasting) in HFD and LFD treated mice for 1 week (N=15/group), 4 weeks (N=15/ group) and 12 weeks (N=10/group). *** P < 0.001, ****P<0.0001. Multiple two-tailed t-test, LFD vs HFD.
c) Fat mass as measured by EchoMRI. ) in HFD and LFD treated mice for 1 week (N=11/group), 4 weeks (N=15/ group) and 12 weeks (N=10/group). ****P<0.0001. Multiple two-tailed t-test, LFD vs HFD.

Intraperitoneal Glucose Tolerance Test (ipGTT, 2g/kg glucose), after 6 hr fast in
d) 1 week LFD vs HFD mice (N= 15/group). Two-way repeated measures ANOVA with post-hoc Sidak's multiple comparison test (*** P< 0.001, , ****P<0.0001).
e) 4 weeks LFD vs HFD mice (N= 15/group). two-way repeated measures ANOVA with post-hoc Sidak's multiple comparison test (*** P< 0.001, , ****P<0.0001).
f) 12 weeks LFD vs HFD mice (N= 10/group). two-way repeated measures ANOVA with post-hoc Sidak's multiple comparison test (**P< 0.01, ***P< 0.001, ****P<0.0001).
g) Area Under the Curve (AUC, 0 min to 120 min) corresponding to ipGTT in HFD and LFD treated mice for 1 week, 4 weeks and 12 weeks , ****P<0.0001. Multiple two-tailed t-test, LFD vs HFD.

Oral Glucose Tolerance Test (oGTT), after 6 hr fast in
h) 1 week LFD vs HFD mice (N= 10/group). two-way repeated measures ANOVA with post-hoc Sidak's multiple comparison test.
i) 4 weeks LFD vs HFD mice (N= 10/group). two-way repeated measures ANOVA with post-hoc Sidak's multiple comparison test (*P< 0.05, , ****P<0.0001).
j) 12 weeks LFD vs HFD mice (N= 10/group). two-way repeated measures ANOVA with post-hoc Sidak's multiple comparison test (**P< 0.05, *** P< 0.001, , ****P<0.0001).
k) AUC (0 min to 120 min) corresponding to oGTT in HFD and LFD treated mice for 1 week, 4 weeks and 12 weeks. Multiple two-tailed t-test, LFD vs HFD ****P<0.0001.

Intraperitoneal Insulin Tolerance Test (ipTT, 0.3 U/kg) after 6 hr fast in
l) 1 week LFD vs HFD mice (N= 15/group). two-way repeated measures ANOVA with post-hoc Sidak's multiple comparison test (** P< 0.01 *** P< 0.001, , ****P<0.0001).
m) 4 weeks LFD vs HFD mice (LFD, N= 19; HFD, N=20). two-way repeated measures ANOVA with post-hoc Sidak's multiple comparison test (** P< 0.01 *** P< 0.001, , ****P<0.0001).
n) 12 weeks LFD vs HFD mice (N= 10/group). two-way repeated measures ANOVA with post-hoc Sidak's multiple comparison test (** P< 0.01, *** P< 0.001, , ****P<0.0001).
o) AUC (0 min to 120 min) corresponding to ipTT in HFD and LFD treated mice for 1 week , 4 weeks and 12 weeks , Multiple two-tailed t-test, LFD vs HFD ****P<0.0001

p) Plasma Insulin levels (6 hours fasting) in HFD and LFD treated mice for 1 week (LFD N=14; HFD N=15), 4 weeks (N=15/ group) and 12 weeks (N=10/group). ** P < 0.01, ****P<0.0001. Multiple two-tailed t-test, LFD vs HFD.
q) Plasma glucagon levels (6 hours fasting) in HFD and LFD treated mice for 1 week (N=15/ group), 4 weeks (N=15/ group) and 12 weeks (N=10/group). * P < 0.05, ****P<0.0001. Multiple two-tailed t-test, LFD vs HFD.
r) β cell volume in HFD and LFD treated mice for 1 week (LFD N=11; HFD N=12) 4 weeks (LFD N=12; HFD N=18) and 12 weeks (LFD N=14; HFD N=15) . * P < 0.05, **. Multiple two-tailed t-test.

All data represented as mean± SEM

### Figure S2

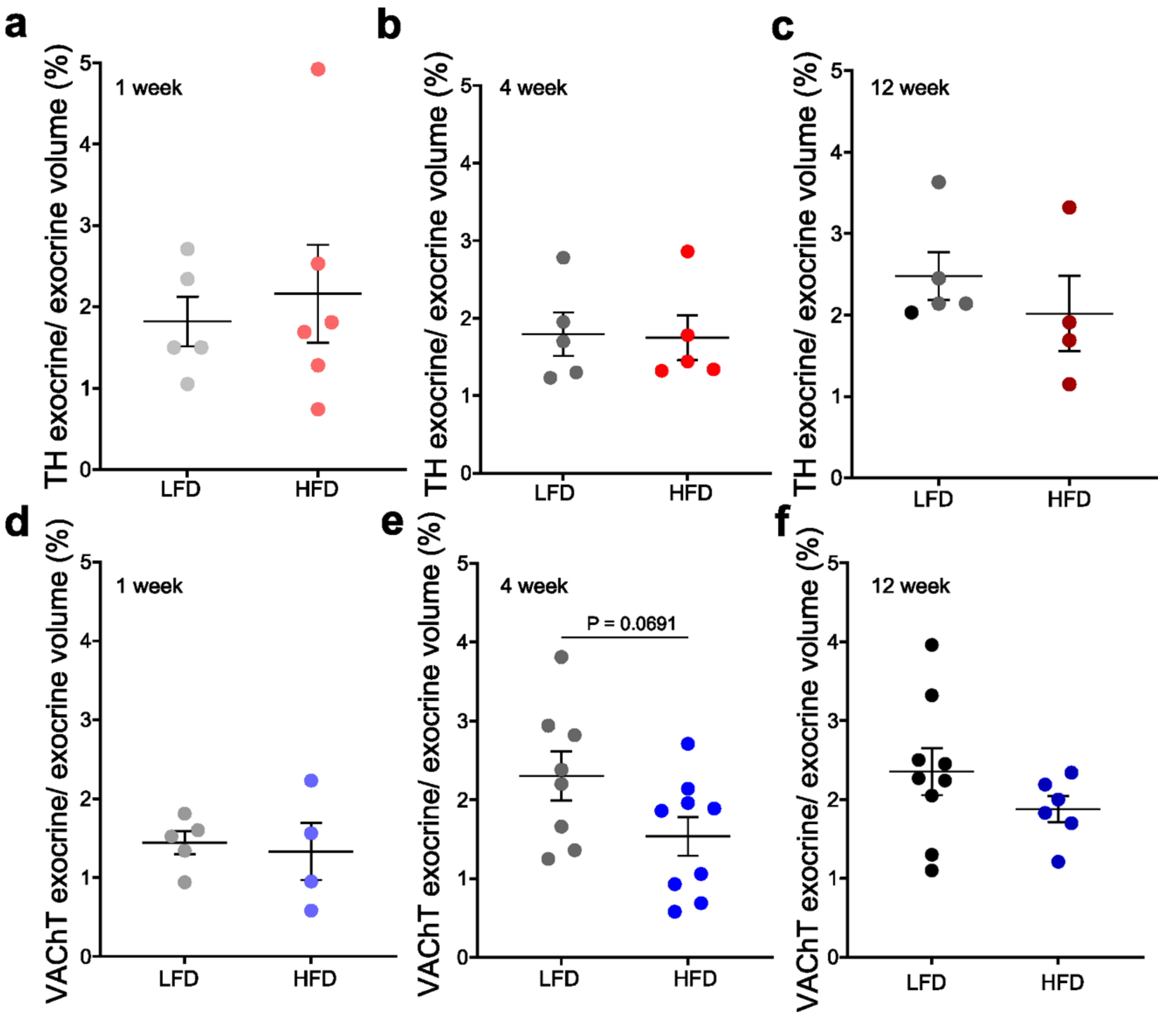


**Figure S2**. **Analysis of exocrine pancreatic autonomic innervation in HFD**.

Quantification of sympathetic TH+ innervation density, as percentage of total pancreas tissue volume in mice receiving HFD or LFD for
a)1 week**,**
b) 4 weeks
c) 12 weeks.
Quantification of parasympathetic VAChT+ innervation density, as percentage of total pancreas tissue volume in mice receiving HFD or LFD for
d)1 week**,**
e) 4 weeks
f) 12 weeks.

All data represented as mean± SEM.

### Figure S3

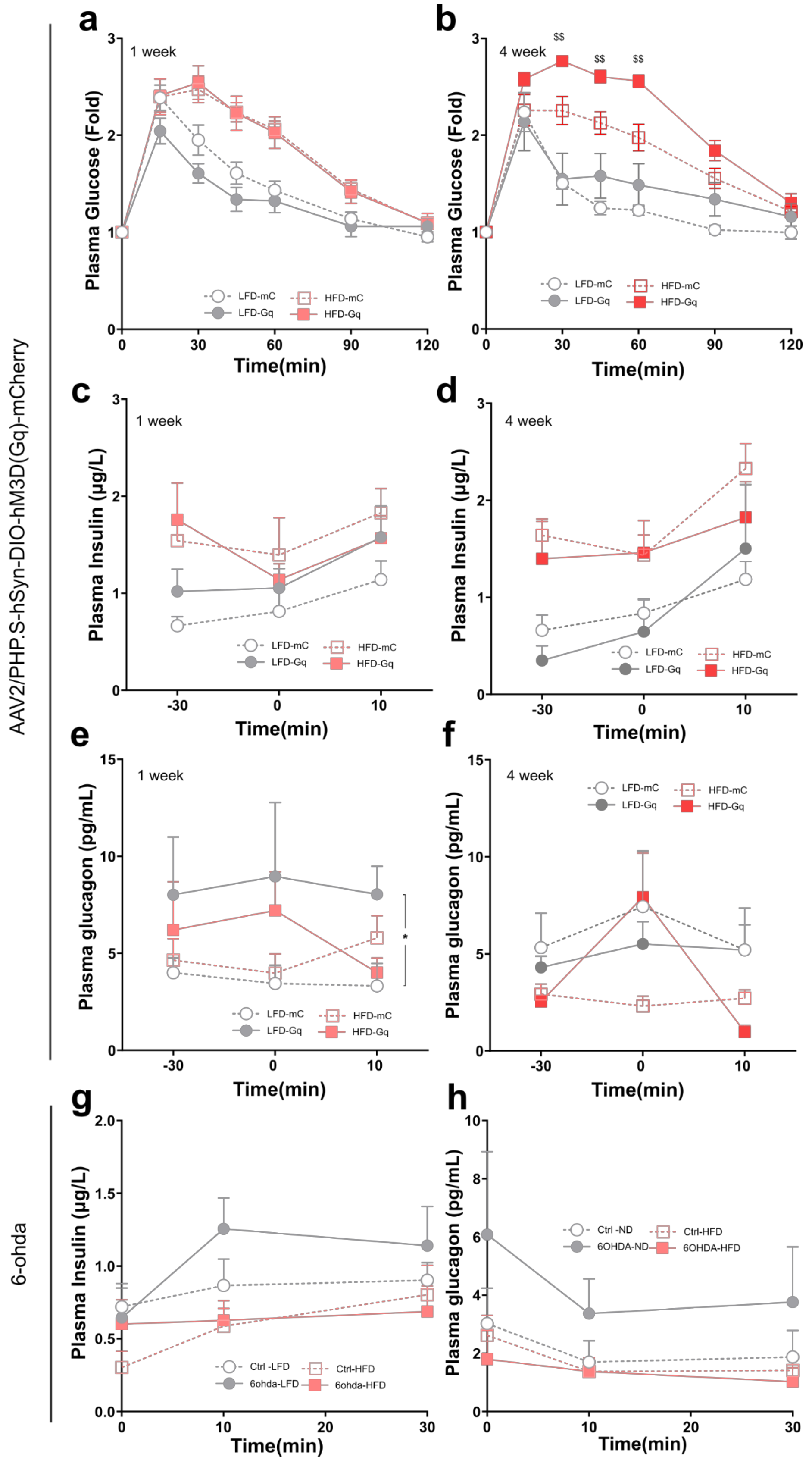

AAV2/PHP.S-hSyn-DIO-hM3D(Gq)-mCherry
6-ohda
a
1 week
Plasma Glucose (Fold)
Time(min)
LFD-mC
LFD-Gq
HFD-mC
HFD-Gq
b
4 week
$$
c
1 week
Plasma Insulin (µg/L)
d
4 week
e
1 week
Plasma glucagon (pg/mL)
*
f
4 week
g
Ctrl -LFD
6ohda-LFD
Ctrl-HFD
6ohda-HFD
h
Ctrl -ND
6OHDA-ND
Ctrl-HFD
6OHDA-HFD

**Figure S3. Effects of pancreatic sympathetic activation and ablation in Pancreatic hormones.**

a) Fold of Glucose Tolerance Test (ipGTT, 2g/kg glucose) in Figure 4d, from the time of glucose administration.
b) Fold of Glucose Tolerance Test (ipGTT, 2g/kg glucose) in Figure 4g, from the time of glucose administration. Two-way ANOVA , with post-hoc Tukey's multiple comparison test, $$ $P<0.01$, HFD-mC vs HFD-Gq

Plasma insulin levels during GTT in at −30, 0 and 10 min in mice receiving intracoeliac injection of AAVphps-hSyn-DIO-hM3D(Gq)-mCherry and intrapancreatic injection of AAV8-hSyn-CRE-eGFP after
c) 1 week of LFD or HFD
d) 4 weeks of LFD or HFD

Plasma glucagon levels during GTT in at −30, 0 and 10 min in mice receiving intracoeliac injection of AAVphps-hSyn-DIO-hM3D(Gq)-mCherry and intrapancreatic injection of AAV8-hSyn-CRE-eGFP after
e) 1 week of LFD or HFD. Mixed-effect analysis with post-hoc Tukey's multiple comparison test. * $P<0.05$ AT 10'.
f) 4 weeks of LFD or HFD

Plasma insulin levels during GTT in at −30, 0 and 10 min in mice treated with intrapancreatic injections of 6-OHDA after
g) 1 week of LFD or HFD
h) 4 weeks of LFD or HFD

All data represented as mean± SEM

### Figure S4

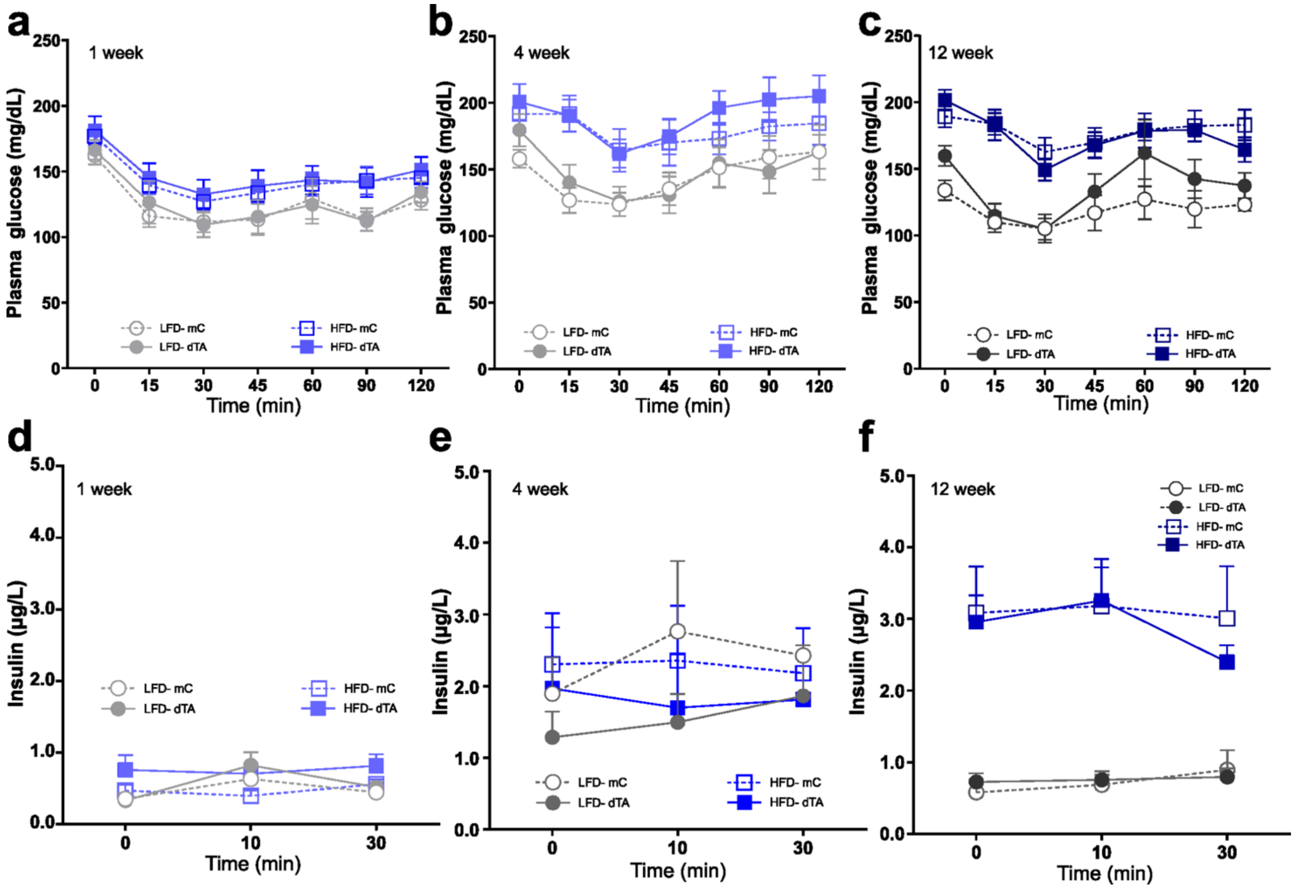


**Figure S4. Effects of pancreatic parasympathetic ablation in glucose regulation and Pancreatic hormones.**

Insulin Tolerance Test (0.3 u/kg, ip) in HFD and LFD treated mice expressing AAV8-EF1a-mCherry-flex-dtA (LFD-dTA and HFD-dTA) or AAV8-EF1a-DIO-mCherry 1 weeks after

a) 1 week of LFD or HFD
b) 4 weeks of LFD or HFD
c) 12 weeks of LFD or HFD

Plasma insulin levels during GTT at −30, 0 and 10 min 1 weeks after

d) 1 week of LFD or HFD
e) 4 weeks of LFD or HFD
f) 12 weeks of LFD or HFD

All data represented as mean± SEM

### Figure S5

a b

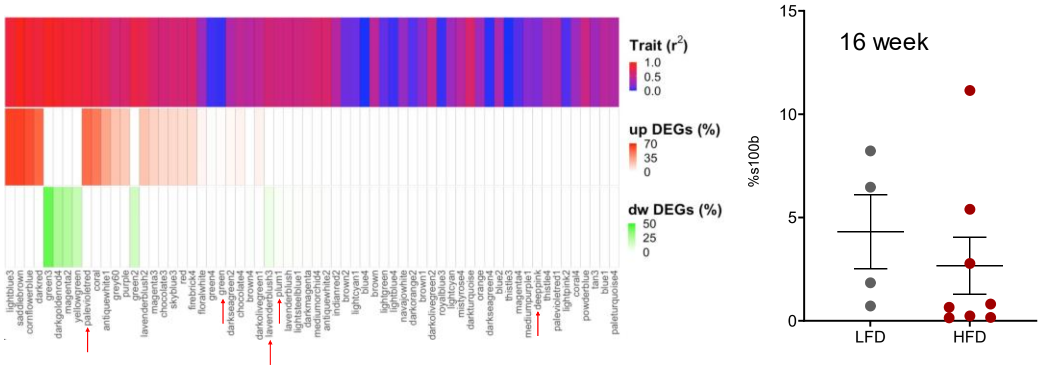


**Figure S5.**

a) Significantly up-regulated and down-regulated differentially expressed genes in each gene module from bulk RNAseq data reanalyzed from reference 39.

b) Quantification of S100b expression within islets as percentage of islet area, in mice receiving HFD for 16 weeks.

All data represented as mean± SEM
